# Ptr1 and ZAR1 immune receptors confer overlapping and distinct bacterial pathogen effector specificities

**DOI:** 10.1101/2022.05.16.492216

**Authors:** Ye Jin Ahn, Haseong Kim, Sera Choi, Carolina Mazo-Molina, Maxim Prokchorchik, Ning Zhang, Boyoung Kim, Hyunggon Mang, Hayeon Yoon, Cécile Segonzac, Gregory B. Martin, Alex Schultink, Kee Hoon Sohn

## Abstract

Nucleotide-binding and leucine-rich repeat receptors (NLRs) detect pathogen effectors inside the plant cell. To identify *Nicotiana benthamiana* NLRs (NbNLRs) with novel effector recognition specificity, we designed an NbNLR VIGS library and conducted a rapid reverse genetic screen. During the NbNLR VIGS library screening, we identified that *N. benthamiana* homolog of Ptr1 (<u>P</u>SEUDOMONAS SYRINGAE PV. <u>T</u>OMATO <u>R</u>ACE <u>1</u> RESISTANCE) recognizes the *Pseudomonas* effectors AvrRpt2, AvrRpm1, and AvrB.

We demonstrated that recognition of the *Xanthomonas* effector AvrBsT and the *Pseudomonas* effector HopZ5 in *N. benthamiana* is conferred independently by *N. benthamiana* homolog of Ptr1 and ZAR1 (HOP<u>Z</u>-<u>A</u>CTIVATED <u>R</u>ESISTANCE <u>1</u>). In addition, we showed that the RLCK XII family protein JIM2 (XOP<u>J</u>4 <u>IM</u>MUNITY <u>2</u>) physically interacts with AvrBsT and HopZ5 and is required for the NbZAR1-dependent recognition of AvrBsT and HopZ5. The recognition of multiple bacterial effectors by Ptr1 and ZAR1 in *N. benthamiana* demonstrates a convergent evolution of effector recognition across plant species. Identification of key components involved in Ptr1 and ZAR1 mediated immunity would reveal unique mechanisms of expanded effector recognition and be useful for engineering resistance in solanaceous crops.

## INTRODUCTION

Pathogen recognition is a key process that leads to plant survival. Plant immune receptors can recognize invading pathogens and activate defense responses. The first level of defense is regulated at the plant cell surface where microbe-associated molecular patterns are recognized by pattern-recognition receptors and activate pattern-triggered immunity (PTI) (Jones & Dangl, 2006). Successful pathogens can suppress PTI with specialized proteins called effectors. Bacterial pathogens often secrete the effectors into a host cell using a syringe-like structure called type III secretion system (T3SS). Once in the plant cell, the effectors target key components of immune signaling, alter cell physiology, and enhance bacterial proliferation in the plant (Büttner, 2016; Ngou *et al*., 2022a).

In a resistant host, effectors are recognized by nucleotide-binding and leucine-rich repeat receptors (NLRs) encoded by disease resistance (*R*) genes (Dangl & Jones, 2001). The recognized effectors are called avirulence (Avr) effectors which activate corresponding NLRs and trigger effector-triggered immunity (ETI) (Jones & Dangl, 2006). ETI is often accompanied by the hypersensitive response (HR), a form of localized programmed cell death (PCD) at the infection site (Jones & Dangl, 2006; Cui *et al*., 2015). NLRs can be categorized according to their N-terminal domain: coiled-coil (CC) NLRs (CNLs), Toll/interleukin-1 receptor/resistance protein (TIR) NLRs (TNLs), and RPW8-like coiled-coil NLRs (RNLs) (Jones *et al*., 2016). Whereas CNLs and TNLs function as effector-recognizing sensors, RNLs play an important role in immune activation and HR development with other key regulators such as <u>E</u>NHANCED <u>D</u>ISEASE <u>S</u>USCEPTIBILITY <u>1</u> (EDS1) (Lapin *et al*., 2020; Huang *et al*., 2022). One of the best-studied sensor NLRs is HOP<u>Z</u>-<u>A</u>CTIVATED <u>R</u>ESISTANCE <u>1</u> (ZAR1) which forms a resistosome upon effector recognition and functions as a calcium channel in the plasma membrane (Wang *et al*., 2019; Bi *et al*., 2021). In contrast, other sensor NLRs require <u>N</u>LR-<u>R</u>EQUIRED FOR <u>C</u>ELL DEATH (NRC) or helper RNLs such as <u>A</u>CTIVATED <u>D</u>ISEASE <u>R</u>ESISTANCE 1 (ADR1), and <u>N</u> <u>R</u>EQUIREMENT <u>G</u>ENE 1 (NRG1) for cell death (Wu *et al*., 2017; Lapin *et al*., 2022). Recently, NRG1 was shown to form a calcium channel and regulate calcium influx which leads to HR (Jacob *et al*., 2021).

Effector recognition by NLRs is diverse, but in many cases NLRs indirectly detect effectors by monitoring host proteins that are targeted by the effectors (Jones *et al*., 2016). Effectors target multiple host proteins to enhance bacterial virulence, but some host targets work as immune sensors and activate NLRs. The effector target protein can be categorized as a guardee or a decoy depending on its involvement in disease susceptibility during infection (Van Der Hoorn & Kamoun, 2008). Some NLRs confer recognition of multiple effectors by guarding a common host target protein. A well-studied host target of effectors in *Arabidopsis* is <u>R</u>PM1-<u>IN</u>TERACTING PROTEIN <u>4</u> (RIN4). RIN4 negatively regulates PTI, but upon bacterial flagellin recognition, RIN4 derepresses immune responses and enhances PTI (Kim *et al*., 2005; Chung *et al*., 2014). Several bacterial effectors induce post-translational modifications on the conserved NOI (nitrate-induced) domain of RIN4 to inhibit immune responses in the host (Toruño *et al*., 2019). Among these effectors, five sequence-unrelated effectors target RIN4 and activate ETI in *Arabidopsis*. Two *Pseudomonas* effectors AvrRpm1 and AvrB induce phosphorylation at Thr166 of RIN4 which activates the CNL protein <u>R</u>ESISTANCE TO <u>P</u>. SYRINGAE PV. <u>M</u>ACULICOLA <u>1</u> (RPM1) (Chung *et al*., 2011; Liu *et al*., 2011; Redditt *et al*., 2019). Similarly, *Pseudomonas* and *Xanthomonas* effectors, HopZ5 and AvrBsT, respectively, acetylate Thr166 and activate RPM1 (Choi *et al*., 2021). Lastly, the *Pseudomonas* effector AvrRpt2 cleaves RIN4 at two major sites leading to the activation of the CNL protein <u>R</u>ESISTANCE TO <u>P</u>. <u>S</u>YRINGAE <u>2</u> (RPS2) in *Arabidopsis* (Axtell & Staskawicz, 2003; Mackey *et al*., 2003; Day *et al*., 2005). Interestingly, AvrRpt2 is recognized across several plant species. In a wild apple species *Malus x robusta 5*, the CNL FB_MR5 is activated by apple RIN4 upon its cleavage by AvrRpt2 from the fire blight pathogen *Erwinia amylovora* (Vogt *et al*., 2013; Prokchorchik *et al*., 2020). In tomato, the CNL *<u>P</u>. syringae* pv. *tomato* race <u>1</u> (Ptr1) protein recognizes AvrRpt2 in race 1 strains of *P. syringae* pv. *tomato* and its *Ralstonia* homolog RipBN (Mazo-Molina *et al*., 2020). Remarkably, while AvrRpt2 is present in *Pseudomonas, Erwinia*, and *Ralstonia* (Innes *et al*., 1993; Zhao *et al*., 2006; Mazo-Molina *et al*., 2019), the corresponding NLRs, RPS2, MR5, and Ptr1, share little sequence homology suggesting that these genes emerged independently. *NLR* genes required for recognition of AvrRpm1, AvrB, HopZ5, and AvrBsT have not been identified in *Solanum* or *Malus* species.

The recognition of several YopJ family bacterial effectors by ZAR1 requires receptor-like cytoplasmic kinases (RLCKs). For instance, ZAR1-mediated recognition of HopZ1a from *P. syringae* in *Arabidopsis thaliana* requires the RLCK XII family protein HOP<u>Z</u> <u>E</u>TI-<u>D</u>EFICIENT <u>1</u> (ZED1) and the RLCK VII family proteins SZE1 and SZE2 (Lewis *et al*., 2010; Lewis *et al*., 2013; Liu *et al*., 2019; Hu *et al*., 2020). In *Nicotiana benthamiana*, XopJ4 from *Xanthomonas perforans* is recognized by ZAR1 with the requirement of XOP<u>J</u>4 <u>IM</u>MUNITY <u>2</u> (JIM2), another RLCK XII family protein (Schultink *et al*., 2019). Furthermore, in association with diverse RLCK proteins, ZAR1 recognizes multiple bacterial effectors that do not belong to the YopJ family (Martel *et al*., 2020). The RLCKs that are required for the NLR function include <u>PB</u>S1-like proteins (PBLs from RLCK group VII) and <u>Z</u>ED1-related <u>k</u>inases (ZRKs from RLCK group XII) (Martel *et al*., 2020). The *Xanthomonas* effector AvrAC uridylates PBL1 which promotes PBL1 interaction with ZRK1 and ZAR1 (Wang *et al*., 2015) and results in the formation of a resistosome (Wang *et al*., 2019; Bi *et al*., 2021). ZAR1 recognition of the *Pseudomonas* effectors HopX1, HopO1, and HopBA1 requires ZED1, ZRK3, and ZRK2, respectively, and HopX1 additionally requires the PBL SZE1 (Laflamme *et al*., 2020; Martel *et al*., 2020). The elaborate network of host sensors enhances effector recognition by NLRs and may be useful for recognizing additional effectors. For example, AvrRpm1 induces ADP-ribosylation not only on RIN4 but also on ten other NOI proteins in *Arabidopsis* (Redditt *et al*., 2019). Similar to PBLs and ZRKs, AvrRpm1 targeted NOI proteins might function as decoys for NLR activation in some plant species.

In this study, we conducted a rapid reverse genetic screen to identify NLRs that recognize *Arabidopsis* RIN4-targeting effectors in *N. benthamiana*. An NbNLR VIGS library was designed to silence 301 *N. benthamiana* NLRs (NbNLRs) using VIGS (virus-induced gene silencing). The screening revealed that the *N. benthamiana* homolog of Ptr1 (NbPtr1) recognizes AvrRpt2, AvrRpm1, AvrB, HopZ5, and AvrBsT. In addition to NbPtr1, we discovered that HopZ5 and AvrBsT are also recognized independently by *N. benthamiana* homolog of ZAR1 (NbZAR1) with the requirement of the RLCK XII family protein, JIM2. Using co-immunoprecipitation, we showed that, unlike XopJ4, HopZ5 and AvrBsT interact strongly with JIM2. Based on these findings, we conclude that NbPtr1 and NbZAR1 show overlapping and distinct effector recognition specificities and provide an opportunity to develop solanaceous crops resistant to multiple bacterial pathogens.

## MATERIALS AND METHODS

### Bacterial strains growth and transformation

*Escherichia coli* DH5α and *Agrobacterium tumefaciens* AGL1 or GV3101 were grown on low-salt Luria-Bertani (LB) liquid or solid agar (20g/L) medium (Duchefa Biochemie, Haarlem, The Netherlands) containing appropriate antibiotics. Plasmids used in this study were transformed to *E. coli, A. tumefaciens*, or *Pseudomonas syringae* pv. *tomato* DC3000 Δ*hopQ1-1* (*Pst* Δ*hopQ1-1*) using electroporation. *Pst* Δ*hopQ1-1* and *Xanthomonas campestris* pv. *vesicatoria* strains Bv5-4a and Ds1.1 were grown in King’s B (KB) media (Duchefa Biochemie, Haarlem, The Netherlands) containing appropriate antibiotics. *Xanthomonas perforans* strain 4B was grown in KB media (1% peptone, 0.15% K_2_HPO_4_, 1.5% (w/v) glycerol, 5 mM MgSO_4_, pH 7.0) containing appropriate antibiotics.

### Plant materials and growth condition

*Nicotiana benthamiana* and *Capsicum annuum* 35001 plants were grown on Gyeongju Chemical Mix soil (70% cocopeat, 13% perlite, 13% peatmoss, and 3% zeolite; Dong-sin, Gyeongju, Korea) in short-day condition (11 hours of light per day) at 22□. *N. benthamiana* accessions used in this study are Nb-0 and Nb-1 (Fig. S1) (Bombarely *et al*., 2012). The difference between Nb-0 and Nb-1 accessions is the ability of the plant to recognize the XopJ4 effector, which occurs in Nb-0 but is absent in Nb-1. This difference may be caused by reduced expression of JIM2 in Nb-1 (Fig. S1) (Schultink *et al*., 2019). Nb-0 was used for the experiments in this study unless Nb-1 is indicated. *Nicotiana glutinosa* and *S. lycopersicoides* introgression line LA4245 were grown on Cornell Osmocote Mix soil (0.16 m^3^ peatmoss, 0.34 m^3^ vermiculite, 2.27 kg lime, 2.27 kg Osmocote Plus 15-9-12 and 0.54 kg Uni Mix 11-5-11; Everris, Geldermalsen, The Netherlands) in long-day condition at 24□ with light and 20□ in the dark. Seeds of *S. lycopersicoides* introgression line LA4245 were obtained from the Tomato Genetics Resource Center (https://tgrc.ucdavis.edu/lycopersicoides_ils.aspx). The genotype of LA4245 can be heterozygous for the presence of *Ptr1* (*Ptr1 ptr1*) (LA4245-R) or homozygous for the lack of the gene (*ptr1 ptr1*) (LA4245-S) (Mazo-Molina *et al*., 2020).

### Plasmid construction and virus-induced gene silencing

To generate VIGS fragments targeting NLRs in *Nicotiana benthamiana*, nucleotide sequences of 307 NbNLRs (Seong *et al*., 2020) were analyzed, and 120 bp-long fragments specific to each of 301 *NLR* genes were selected (Table S1). Four to six VIGS-fragments were grouped to make one VIGS-combination (VIGS-com) (Table S1). In total, 55 VIGS-com were designed. DNA fragments of VIGS-com flanked by BsaI sites were synthesized (Twist Biosciences, USA) and cloned into the pUC19b vector (Jayaraman *et al*., 2017). Subsequently, VIGS-com were assembled into a Golden Gate compatible pTRV2-66 (Choi *et al*., 2021). All VIGS constructs in addition to VIGS-com cloned in this study were assembled into pTRV2-66 (Choi *et al*., 2021).

To silence target NbNLRs, *A. tumefaciens* AGL1 or GV3101 transformed with pYL192 and pTRV2-66 carrying VIGS-com were grown on low-salt LB media at 28□ overnight. Overnight cultures were resuspended in VIGS buffer (10 mM MgC1_2_, 1 mM MES (2-(*N*-morpholino)ethanesulfonic acid) [pH 5.6]) and diluted to the OD_600_ of 0.5. *A. tumefaciens* transformed with pYL192 were co-infiltrated with *A. tumefaciens* transformed with pTRV2-66 carrying VIGS construct into leaves of *N. benthamiana* grown in short-day condition for three to five days after transplantation. Subsequently, silenced *N. benthamiana* plants were further grown for four to five weeks before effector-induced PCD was tested. For VIGS in *Capsicum annuum*, the experimental procedure was identical to *N. benthamiana* except that the plants were kept in a short-day condition at 18□ and 50% humidity for seven days after *A. tumefaciens* infiltration.

### *Agrobacterium*-mediated transient transformation and ion-leakage assays

Overnight cultures of *A. tumefaciens* were resuspended in an infiltration buffer (10 mM MgCl_2_, 10mM MES [pH 5.6]) and diluted to the OD_600_ values indicated in the figure legends. Five-week-old *N. benthamiana* and *N. glutinosa* were infiltrated with *A. tumefaciens* AGL1 and GV3101, respectively using a needleless syringe. PCD was measured at two to three days post-infection (dpi). To quantify PCD, four replicates of plant samples (three leaf discs for each replicate, 8 mm in diameter) were collected at 2 dpi and floated on 2 ml of sterile water in a multi-well culture plate. Conductivity was measured using EC33 (Horiba, Kyoto, Japan) after two hours of incubation.

### Bacterial growth assay

For the bacterial growth assays with *Pst* Δ*hopQ1-1* strains, the bacteria were grown on KB media at 28°C for two days. *Pst* Δ*hopQ1-1* strains were resuspended in 10 mM MgCl_2_ at a final concentration of 5 × 10^4^ CFU/ml (OD_600_ = 0.0001) and infiltrated to five-week-old *N. benthamiana* leaves. At 4 dpi, four replicates of plant samples (two leaf discs for each replicate, 8mm in diameter) were ground in sterile 10 mM MgCl_2_, serially diluted, and plated on KB plates with appropriate antibiotics. The number of bacterial colonies was counted from the plates after two days of incubation at 28°C. For the bacterial growth assays with *Xanthomonas perforans*, the bacteria were grown overnight in liquid KB media at 30°C with appropriate antibiotics, washed, and diluted to OD_600_ = 0.0001 with 10 mM MgCl2 for infiltration into leaf tissue. At 6 dpi, four leaf punches were collected for each replicate, the punches were homogenized in sterile water, and the solution was serially diluted and plated on KB plates. The colonies were counted after two days of incubation at 30°C.

### Immunoblot and co-immunoprecipitation analyses

For total protein extraction, nine leaf discs (8 mm in diameter) were collected from five-week-old *N. benthamiana* leaves after infiltration of *A. tumefaciens*. Leaf samples were ground with a pestle and 200 μl of SDS protein loading buffer (250 mM Tris-HCl, 8% SDS, 40% glycerol, 100 mM DTT, 0.1% bromophenol blue) was added. Total extracts were then boiled at 95□ for 10 minutes. Fifteen μl of total extracts from each sample was loaded for protein electrophoresis.

For co-immunoprecipitation, 0.5 g of agroinfiltrated leaf tissue was collected and grounded with a liquid nitrogen-chilled mortar and a pestle. Six ml of protein extraction buffer (1 tablet of Sigma Complete™ mini protease inhibitor, IGEPAL 0.2%, 10mM DTT, PVPP 2%, GTEN buffer) was added. Protein extracts were then centrifuged at 2,600 g at 4□ for 10 minutes. The supernatant was filtered through Merck MiraCloth (Merk, Kenilworth, USA). Twenty μl of anti-GFP beads was mixed with 1.4 ml of filtered extracts (Chromotek, Munich, Germany) and incubated at 4□ for 2 hours. The beads were washed with GTEN buffer and 60 μl of SDS protein loading buffer was added. Subsequently, beads were boiled at 95□ for 10 minutes and 15 μl of protein samples were loaded for protein electrophoresis.

For SDS-PAGE and protein detection, protein samples were loaded in 8 % polyacrylamide gel and subjected to electrophoresis at 130 V for 1 hour. Proteins were transferred to PVDF membranes and incubated in blocking buffer (5% skim milk dissolved in TBST buffer) for 20 minutes. Anti-FLAG antibodies (Sigma-Aldrich, St. Louis, USA; 5,000 times dilution), anti-MYC antibodies (Cell Signaling Technology, Danvers, USA; 4,000 times dilution), anti-HA antibodies (Sigma-Aldrich, St. Louis, USA; 2,000 times dilution), or anti-GFP antibodies (Invitrogen, Waltham, USA; 4,000 times dilution) were then added and incubated for 1 hour. Membranes were washed with TBST buffer (three times, 7 minutes each) and incubated with anti-mouse-HRP or anti-rabbit-HRP (Sigma-Aldrich; 20,000 times dilution) for 1 hour. Subsequently, membranes were washed with TBST buffer and the chemiluminescence of the protein samples was visualized using Super Signal West Femto reagents (Thermo Scientific, Waltham, USA) through Image Quant LAS500 (GE Healthcare, Chicago, USA).

## RESULTS

### Construction and screening of the NbNLR VIGS library

To rapidly identify *NLR* genes that confer novel effector recognition specificities, we used VIGS to silence NbNLRs based on the previously annotated 307 NbNLRs (Table S1) (Seong *et al*., 2020). We designed VIGS-fragments that each can silence one NbNLR and screened for reduction in effector-induced programmed cell death (PCD) in the silenced plants (Fig. 1a). To reduce the number of constructs used in the screening, four to six VIGS fragments were assembled into a pTRV2-GG vector (for details, see Materials and Methods). As a result, we created an NbNLR VIGS library that consisted of 55 VIGS constructs targeting 301 NbNLRs (Fig. 1a). Six NLRs were excluded from the library due to short amino acid sequences, high sequence similarities with other NLRs in the same phylogenetic groups, or retardation of growth post-silencing (Table S1) (Adachi *et al*., 2021). The NbNLR VIGS constructs were labeled from Com-1 to Com-55 and transformed to *A. tumefaciens* for VIGS. Subsequently, effector-induced PCD was tested on fully expanded leaves of VIGSed *N. benthamiana* plants (Fig. 1a).

**Fig. 1.**
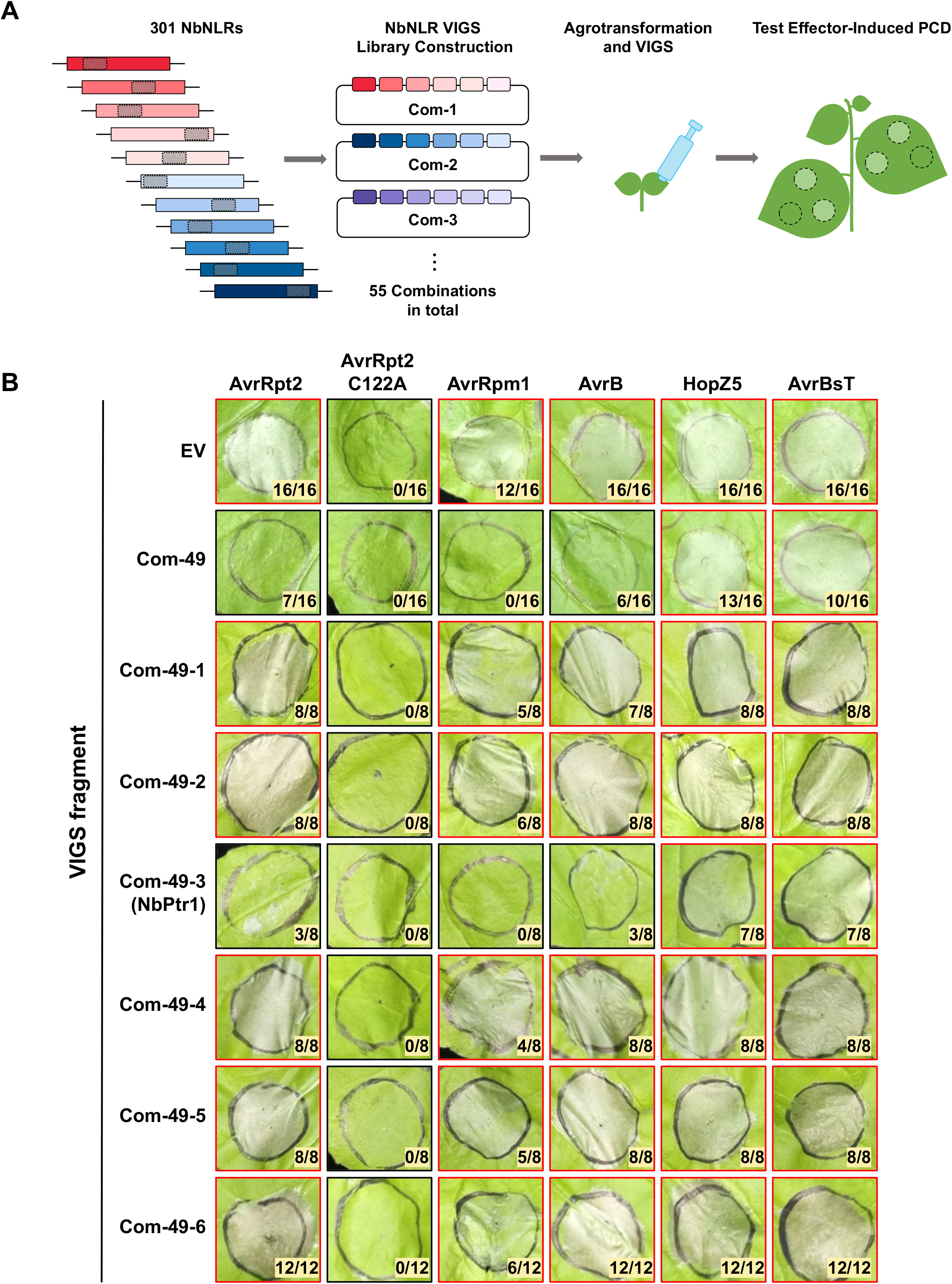
Com-49 VIGS reduces PCD induced by AvrRpt2, AvrRpm1, and AvrB. (a) Schematics of the NbNLR VIGS library construction and screening. 301 annotated *NbNLRs* were silenced using VIGS. Gray boxes on *NbNLRs* indicate VIGS-fragments that are unique to each *NLR*. Four to six VIGS-fragments were assembled into a pTRV2-GG resulting in 55 VIGS-Com constructs. *Agrobacterium* carrying each VIGS-Com was infiltrated into *N. benthamiana* seedlings three to five days after transplantation. The effectors were expressed via agroinfiltration in leaves of mature VIGSed *N. benthamiana*. (b) The *NLR* targeted by Com-49-3 is required for PCD induced by AvrRpt2, AvrRpm1, or AvrB. Com-49 targets six *NLRs*, and Com-49-1 to Com-49-6 were designed to silence each *NLR* separately. The effectors were agroinfiltrated at an OD_600_ of 0.4. The numerator indicates the number of spots with cell death, and the denominator indicates the total number of infiltrations. If PCD count is higher than 50% of total infiltrations, red borders were marked around the photograph, and black borders were used for lower PCD counts. Photographs are representative PCD observed at 2 dpi.

### Silencing of *NbPtr1* significantly reduces AvrRpt2, AvrRpm1, and AvrB-induced PCD

The bacterial T3SS effectors AvrRpt2, AvrRpm1, AvrB, HopZ5, and AvrBsT trigger immune responses in *N. benthamiana* (Staskawicz *et al*., 1987; Minsavage, 1990; Innes *et al*., 1993; Ritter & Dangl, 1995; Jayaraman *et al*., 2017). We used our NbNLR VIGS library to identify the NLR required for the recognition of avirulence effectors (Fig. 1b). To identify the corresponding NLR(s) in *N. benthamiana*, the avirulence effectors were expressed via *Agrobacterium*-mediated transient transformation (hereafter, agroinfiltration) in *N. benthamiana* silenced with NbNLRs (Fig. 1b). In EV-VIGS control plants, AvrRpt2, AvrRpm1, AvrB, HopZ5, and AvrBsT induced a strong PCD whereas AvrRpt2 C122A, an inactive variant of AvrRpt2, did not (Axtell *et al*., 2003) (Fig 1b). Among the plants silenced with the NbNLR VIGS constructs, Com-49 VIGS plants showed a significant reduction of the PCD induced by AvrRpt2, AvrRpm1, and AvrB (Fig. 1b). Since Com-49 VIGS construct targets six different NbNLRs (Table S1), an additional VIGS experiment was conducted to silence each of the individual NbNLRs targeted by Com-49 VIGS. Interestingly, only Com-49-3 VIGS plants showed a loss of PCD induced by AvrRpt2, AvrRpm1, and AvrB suggesting that the NLR targeted by Com-49-3 VIGS can recognize these avirulence effectors (Fig. 1b). In contrast, Com-49-3 VIGS did not significantly affect HopZ5 or AvrBsT-induced PCD (Fig. 1b). Interestingly, the NLR targeted by Com-49-3 VIGS corresponds to the *N. benthamiana* homolog of Ptr1 (NbPtr1) which was recently shown to recognize *P. syringae* and *R. solanacearum* effectors AvrRpt2 and RipBN, respectively (Mazo-Molina *et al*., 2020).

### NbPtr1 recognizes AvrRpt2, AvrRpm1, and AvrB

To increase the efficiency of *NbPtr1* silencing, another VIGS construct that targets *NbPtr1* gene was designed. The new *NbPtr1*-silencing VIGS construct (NbPtr1-VIGS) targets the first 300 bp of *NbPtr1* gene that encompasses Com-49-3 VIGS target region (Fig. S2). PCD induced by agroinfiltration of AvrRpt2, AvrRpm1, and AvrB was completely abolished in NbPtr1-VIGS plants (Fig. 2a). Interestingly, we consistently observed that HopZ5 or AvrBsT-induced PCD was partially reduced in NbPtr1-VIGS compared to EV-VIGS plants (Fig. 2a). Next, we quantified NbPtr1 transcript accumulation using qRT-PCR in EV-VIGS, Com-49-3 VIGS, and NbPtr1-VIGS plants (Fig. 2b). The expression of *NbPtr1* was further reduced in NbPtr1-VIGS plants compared to Com-49-3 VIGS plants (Fig. 2b). Therefore, we chose NbPtr1-VIGS to silence *NbPtr1* for the subsequent experiments.

**Fig. 2.**
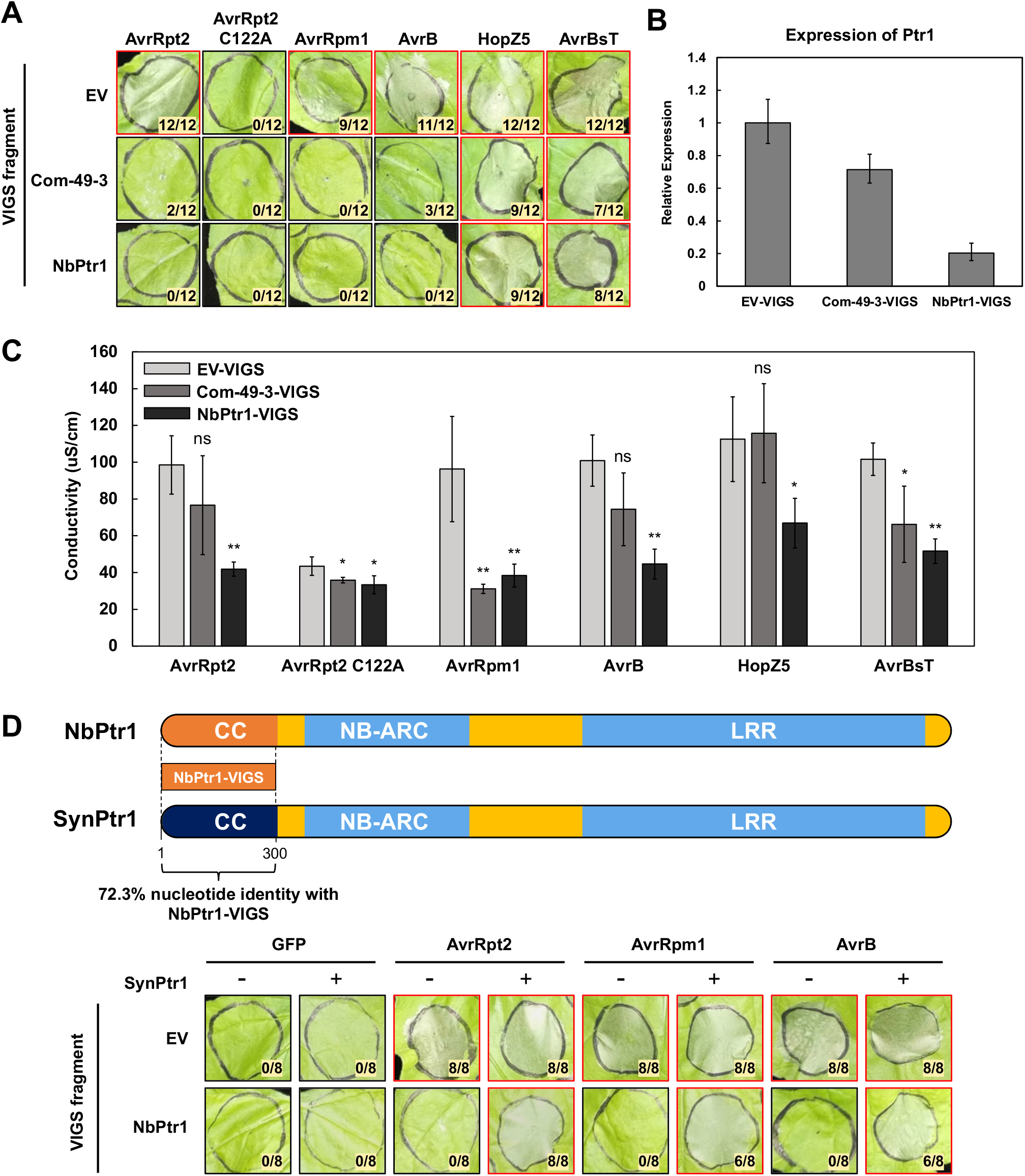
NbPtr1 recognizes AvrRpt2, AvrRpm1, and AvrB. (a) NbPtr1-VIGS construct improves NbPtr1 silencing efficiency. AvrRpt2, AvrRpt2 C122A, AvrRpm1, AvrB, HopZ5, and AvrBsT were agroinfiltrated at an OD_600_ of 0.4. The numerator indicates the number of spots with cell death, and the denominator indicates the total number of infiltrations. If PCD count is higher than 50% of total infiltrations, red borders were marked around the photograph, and black borders were used for lower PCD counts. Photographs are representative PCD observed at 2 dpi. (b) NbPtr1 expression level in leaf tissue, measured by quantitative RT-PCR, is reduced in Com49 and NbPtr1-VIGSed *N. benthamiana. NbPtr1* expression is normalized to endogenous *NbActin* expression, and *NbPtr1* expression in Com-49-3 VIGS and NbPtr1-VIGS plants is relative to *NbPtr1* expression in EV-VIGSed plants. The data shown are the mean of six technical replicates ± SEM. (c) The conductivity of effector-induced PCD is reduced in NbPtr1-VIGSed plants. AvrRpt2, AvrRpt2 C122A, AvrRpm1, AvrB, HopZ5, and AvrBsT were agroinfiltrated at an OD_600_ of 0.4 using *Agrobacterium* in *N. benthamiana*. Leaf discs were collected at 2 dpi for the conductivity assay. Asterisks indicate a significant difference, estimated by the student’s t-test, effector induced conductivity between EV-VIGS and Com-49-3 VIGS or NbPtr1-VIGS plants; *(*P* < 0.05) and **(*P* < 0.01). ‘ns’ indicates that there is no significant difference (*P* > 0.05). The experiment was conducted two times with similar results. (d) SynPtr1 restores AvrRpt2, AvrRpm1, and AvrB induced PCD in NbPtr1-VIGSed *N. benthamiana*. GFP, AvrRpt2, AvrRpm1, and AvrB were transiently expressed at OD_600_ = 0.4 and SynPtr1 at OD_600_ = 0.05 using *Agrobacterium*. ‘+’ and ‘-’ indicate the presence or the absence of SynPtr1, respectively. The numerator indicates the number of spots with cell death, and the denominator indicates the total number of infiltrations. If PCD count is higher than 50% of total infiltrations, red borders were marked around the photograph, and black borders were used for lower PCD counts. Photographs are representative HR observed at 2 dpi.

To quantify effector-induced PCD in the silenced plants, conductivity measurements of ion leakage were performed (Fig. 2c). The expression of AvrRpm1 showed reduced ion leakage in both Com-49-3 VIGS and NbPtr1-VIGS plants (Fig. 2c). Although the reduction of ion leakage induced by AvrRpt2 and AvrB was not significant in Com-49-3 VIGS plants, the reduction was significant in NbPtr1-VIGS plants (Fig. 2c). For HopZ5 and AvrBsT, the reduction of PCD was only partially visible (Fig. 2a), but there was a reduction of ion leakage in Com-49-3 VIGS plants, and this was more significant in NbPtr1-VIGS plants (Fig. 2c). The ion leakage results suggested that NbPtr1 may recognize HopZ5 and AvrBsT in addition to AvrRpt2, AvrRpm1, and AvrB. In *Nicotiana glutinosa*, the expression of AvrRpt2 does not cause PCD due to lack of a functional Ptr1 (Mazo-Molina *et al*., 2020), so we coexpressed the effectors with Ptr1 from *Solanum lycopersicoides* in *N. glutinosa* to test Ptr1 recognition of the effectors (Fig. 3S). Ptr1 was previously shown to be autoactive and cause PCD when expressed at a high level in *N. glutinosa* (Mazo-Molina *et al*., 2020), so a low OD of agrobacterium carrying Ptr1 was used for the experiment (Fig. 3S). Agroinfiltration of Ptr1 with AvrRpt2, AvrRpm1, AvrB, HopZ5, or AvrBsT induced a strong PCD in *N. glutinosa* (Fig. S3). These results are consistent with *N. benthamiana* VIGS results (Fig. 2a and 2c) and collectively imply that Ptr1 can recognize AvrRpt2, AvrRpm1, AvrB, HopZ5, and AvrBsT. However, we hypothesized that an additional NLR(s) might independently recognize HopZ5 and AvrBsT in *N. benthamiana* since the PCD induced by the expression of these effectors was only partially reduced in NbPtr1-VIGS plants.

**Fig. 3.**
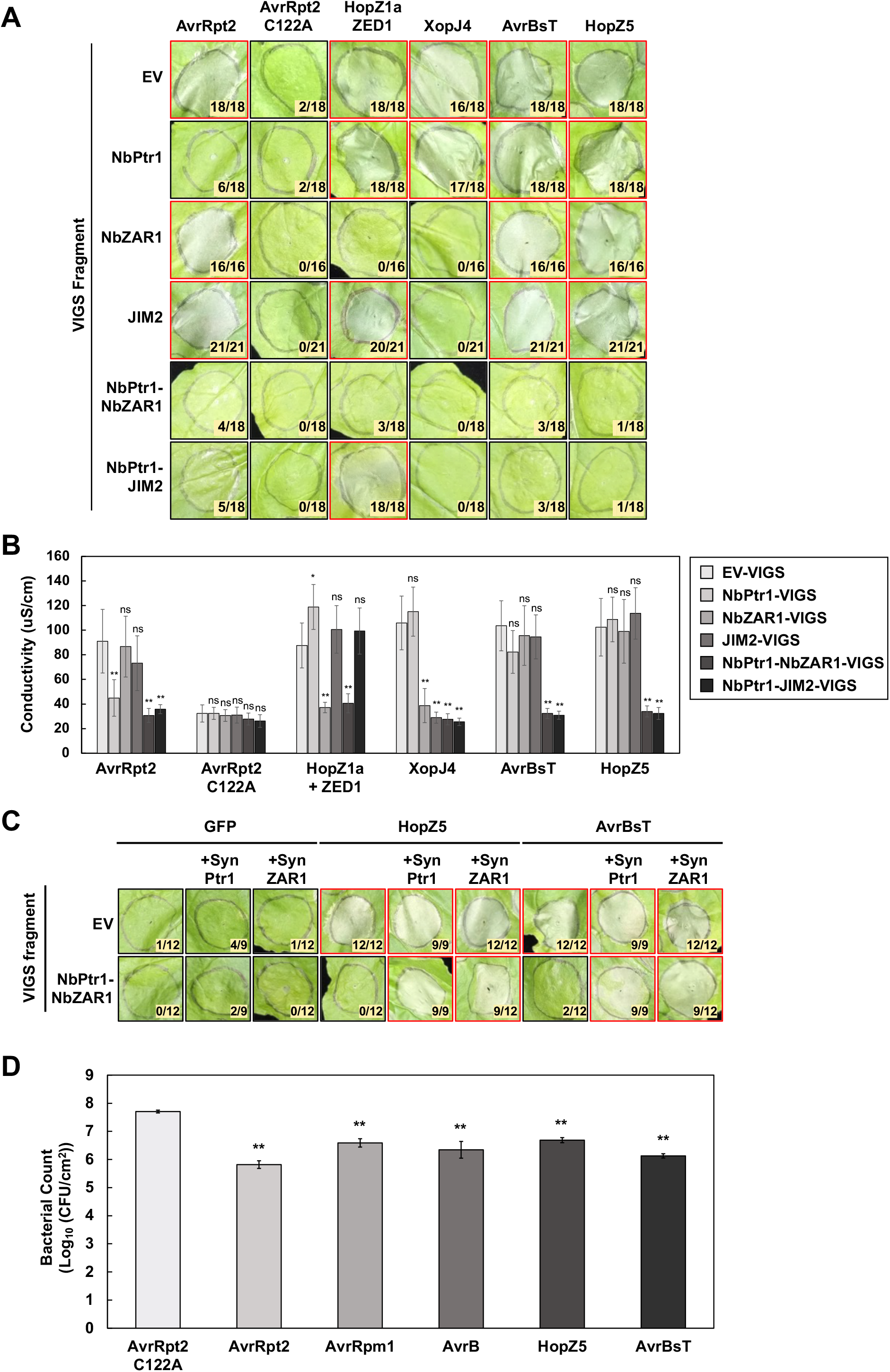
AvrBsT- and HopZ5-triggered immunity is mediated by NbPtr1 and NbZAR1. (a) AvrRpt2, AvrRpt2 C122A, HopZ1a, ZED1, XopJ4, AvrBsT, and HopZ5 were agroinfiltrated at OD_600_ of 0.4 in VIGSed *N. benthamiana*. The numerator indicates the number of spots with cell death, and the denominator indicates the total number of infiltrations. If PCD count is higher than 50% of total infiltrations, red borders were marked around the photograph, and black borders were used for lower PCD counts. Photographs are representative PCD observed at 2 dpi. (b) The conductivity of AvrBsT and HopZ5 induced PCD is reduced in NbPtr1-NbZAR1 or NbPtr1-JIM2-VIGS plants. AvrRpt2, AvrRpt2 C122A, HopZ1a, ZED1, XopJ4, AvrBsT, and HopZ5 were agroinfiltrated OD_600_ of 0.4. Leaf discs were collected at 2 dpi for the conductivity assay. Asterisks indicate significant difference, estimated by student’s t-test, effector induced conductivity between EV-VIGS and NbPtr1-VIGS, NbZAR1-VIGS, JIM2-VIGS, NbPtr1-NbZAR1-VIGS, or NbPtr1-JIM2-VIGS plants; *(*P* < 0.05) and **(*P* < 0.01). ‘ns’ indicates that there is no significant difference (*P* > 0.05). The experiment was conducted four times with similar results. (c) Co-infiltration of SynPtr1 or SynZAR1 restores HopZ5- and AvrBsT-induced PCD in NbPtr1-NbZAR1-VIGS *N. benthamiana*. SynPtr1 and SynZAR1 were designed with alternate codons to evade silencing in NbPtr1-NbZAR1-VIGS plants. GFP, HopZ5, AvrBsT, and SynZAR1 were transiently expressed at OD_600_ = 0.4 and SynPtr1 at OD_600_ = 0.05 using *Agrobacterium*. The numerator indicates the number of spots with cell death, and the denominator indicates the total number of infiltrations. If PCD count is higher than 50% of total infiltrations, red borders were marked around the photograph, and black borders were used for lower PCD counts. Photographs are representative PCD observed at 2 dpi. (d) Delivery of the avirulence effectors confers bacterial growth restriction in wildtype *N. benthamiana*. The growth of *P. syringae* pv. *tomato* DC3000 Δ*hopQ1-1* carrying *avrRpt2 C122A, avrRpt2, avrRpm1, avrB, hopZ5, and avrBsT* at OD_600_ = 0.0001 (5×10^4^ CFU/ml) was quantified at 4 dpi. The data shown are the mean of four biological replicates ± SEM. Asterisks indicate significant difference, estimated by student’s t-test, between bacterial count of the *Pst* Δ*hopQ1-1* carrying *avrRpt2 C122A* strain and the strains carrying *avrRpt2, avrRpm1, avrB, hopZ5*, or *avrBsT*; *(*P* < 0.05) and **(*P* < 0.01). ‘ns’ indicates that there is no significant difference (*P* > 0.05). All experiments were conducted at least two times with similar results.

We conducted a complementation assay of the NbPtr1-VIGS plants with a silencing-proof synthetic *NbPtr1* construct (*SynPtr1*) to demonstrate NbPtr1 requirement for the recognition of AvrRpt2, AvrRpm1, and AvrB (Fig. 2d). *SynPtr1* construct was created with altered codons to avoid silencing in NbPtr1-VIGS plants but encodes the same amino acid as NbPtr1 (Fig. S4). Due to Ptr1 autoactivation (Mazo-Molina *et al*., 2020), a low level of SynPtr1 was used for the complementation assay in *N. benthamiana* (Fig. 2d). Agroinfiltration of SynPtr1 with GFP in EV-VIGS and NbPtr1-VIGS plants did not induce PCD (Fig. 2d). Agroinfiltration of AvrRpt2, AvrRpm1, and AvrB with or without SynPtr1 induced a robust PCD in EV-VIGS plants (Fig. 2d). In NbPtr1-VIGS plants, PCD was restored only when SynPtr1 was coexpressed with AvrRpt2, AvrRpm1, or AvrB (Fig. 2d). These results further demonstrate that NbPtr1 is sufficient for the recognition of AvrRpt2, AvrRpm1, and AvrB.

### AvrBsT and HopZ5 are independently recognized by NbPtr1 and NbZAR1

The partially compromised conductivity level induced by AvrBsT or HopZ5 in NbPtr1-VIGS plants (Fig. 2c) suggests the existence of an additional NLR(s) that recognize these effectors besides NbPtr1 in *N. benthamiana*. Since NbZAR1 was shown to recognize YopJ family effectors HopZ1a and XopJ4 (Baudin *et al*., 2017; Schultink *et al*., 2019), we set out to investigate if NbZAR1 is required for AvrBsT or HopZ5-induced PCD. We silenced *NbZAR1* gene using VIGS and tested for effector-induced PCD (Fig. 3a). As controls, AvrRpt2 and AvrRpt2 C122A were expressed to test for effector recognition by NbPtr1, HopZ1a and ZED1 for NbZAR1, and XopJ4 for NbZAR1 and JIM2 (Fig. 3a). In EV-VIGS plants, all the effectors except AvrRpt2 C122A induced PCD. In NbPtr1-VIGS plants, AvrRpt2-induced PCD was lost due to a lack of NbPtr1 activity. In NbZAR1-VIGS plants, PCD induced by XopJ4 or coexpression of HopZ1a and ZED1 was lost due to a lack of NbZAR1 activity. Importantly, AvrBsT- and HopZ5-induced PCD was not abolished in NbPtr1-VIGS nor NbZAR1-VIGS plants. Since NbPtr1 might independently recognize HopZ5 and AvrBsT in NbZAR1-VIGS plants, this led us to silence both NbPtr1 and NbZAR1 (using an NbPtr1-NbZAR1-VIGS construct). In NbPtr1-NbZAR1-VIGS plants, AvrBsT- and HopZ5-induced PCD was significantly reduced compared to the PCD induced in NbPtr1-VIGS or NbZAR1-VIGS plants. This indicates that HopZ5 and AvrBsT are recognized by NbPtr1 and NbZAR1. Since JIM2 is yet the only *N. benthamiana* RLCK XII family protein identified to be required by NbZAR1 (Schultink *et al*., 2019), we tested for JIM2 requirement in AvrBsT- or HopZ5-triggered immunity using VIGS. In JIM2-VIGS plants, the PCD induced by XopJ4 was abolished, yet AvrBsT or HopZ5 induced strong PCD. However, in NbPtr1-JIM2-VIGS plants, we observed a significant reduction of PCD caused by AvrBsT or HopZ5 that was comparable to what was observed in NbPtr1-NbZAR1-VIGS plants. These results demonstrate that JIM2 is required for AvrBsT- and HopZ5-induced NbZAR1-dependent PCD.

To quantify the PCD results of Fig. 3a, conductivity was measured from the agroinfiltrated leaf samples (Fig. 3b). The expression of AvrBsT or HopZ5 led to a high conductivity in EV-VIGS, NbPtr1-VIGS, NbZAR1-VIGS, and JIM2-VIGS plants (Fig. 3b). However, the conductivity induced by AvrBsT or HopZ5 decreased significantly in NbPtr1-NbZAR1-VIGS or NbPtr1-JIM2-VIGS plants. This result further supports that NbPtr1 and NbZAR1 independently recognize AvrBsT and HopZ5. To confirm this, VIGS-resistant SynPtr1 and synthetic NbZAR1 (SynZAR1) (Fig. S5) were used for a complementation assay in NbPtr1-NbZAR1-VIGS plants (Fig. 3c). Agroinfiltration of SynPtr1 or SynZAR1 with GFP did not induce PCD in EV-VIGS and NbPtr1-NbZAR1-VIGS plants. In EV-VIGS plants, agroinfiltration of HopZ5 or AvrBsT induced PCD regardless of the presence of SynPtr1 or SynZAR1. In NbPtr1-NbZAR1-VIGS plants, HopZ5- and AvrBsT-induced PCD was restored only when the effectors were coexpressed with SynPtr1 or SynZAR1. These results strongly demonstrate that NbPtr1 and NbZAR1 independently recognize AvrBsT and HopZ5.

Based on our results (Fig. 2 and 3), NbPtr1 is activated upon the recognition of AvrRpt2, AvrRpm1, AvrB, HopZ5, and AvrBsT. Since the effectors that are recognized by NbPtr1 are previously reported to modify a guardee called RIN4 in *A. thaliana* (Axtell & Staskawicz, 2003; Mackey *et al*., 2003; Day *et al*., 2005; Chung *et al*., 2011; Liu *et al*., 2011; Redditt *et al*., 2019; Mazo-Molina *et al*., 2020; Choi *et al*., 2021), we tested if *N. benthamiana* RIN4 homologs are required for the effector-induced PCD (Fig. S6). All three *N. benthamiana* RIN4 homologs (Prokchorchik *et al*., 2020) were silenced using NbRIN4-1,2,3 VIGS construct, and we tested for effector-induced PCD. In NbPtr1-VIGS plants, expression of AvrRpt2, AvrB, and AvrRpm1 did not induce PCD, and HopZ5 and AvrBsT expression showed a reduced PCD (Fig. S). In NbRIN4-1,2,3-VIGS plants, expression of the effectors induced PCD except for AvrRpt2 C122A, and this was comparable to what was observed in EV-VIGS plants (Fig. S6). Silencing NbRIN4 homologs did not affect the effector recognition in *N. benthamiana*. This result shows that other host proteins besides NbRIN4 homologs might be required for NbPtr1-mediated immunity.

To test if the recognition of the effectors restricts the growth of a virulent pathogen in *N. benthamiana*, we used *P. syringae* pv. *tomato* DC3000 lacking *hopQ1-1* (*Pst* Δ*hopQ1-1*) as a more natural means of effector delivery (for details, see Methods S2). The effector HopQ1-1 from *P. syringae* pv. *tomato* DC3000 is recognized by the NLR Roq1 and triggers immune responses in *N. benthamiana* (Wei et al., 2007; Schultink et al., 2017). In accordance, the *Pst* Δ*hopQ1-1* strain evades recognition and can cause disease. *Pst* Δ*hopQ1-1* carrying *avrRpt2 C122A, avrRpt2, avrRpm1, avrB, hopZ5*, or *avrBsT* (Choi *et al*., 2018) were infiltrated to wild type *N. benthamiana* for bacterial growth assay (Fig. 3d). *Pst* Δ*hopQ1-1* carrying the avirulence effectors grew to a significantly lower level compared to the strain carrying *avrRpt2 C122A*. The growth restriction of the *Pst* Δ*hopQ1-1* strains carrying the avirulence effectors indicates that NbPtr1 and NbZAR1 can confer disease resistance in *N. benthamiana*.

### HopZ5 and AvrBsT physically associate with JIM2

Since JIM2 and NbZAR1 are required for HopZ5 or AvrBsT-induced PCD, we hypothesized that the avirulence effectors physically associate with JIM2. To test this, we performed co-immunoprecipitation (co-IP) of transiently expressed proteins in *N. benthamiana* (Fig. 4). The C-terminally YFP-tagged XopJ4, AvrRpm1, AvrB, HopZ5, and AvrBsT proteins were coexpressed with C-terminally MYC-tagged JIM2 (JIM2-MYC) in *N. benthamiana* leaf cells using agroinfiltration and enriched from total protein extracts using GFP-trap beads. JIM2-MYC was not co-immunoprecipitated in the samples expressing EV, XopJ4-YFP, AvrRpm1-YFP, or AvrB-YFP. This result is consistent with similar co-immunoprecipitation assays reporting that XopJ4 did not interact with JIM2 (Schultink *et al*., 2019). However, JIM2-MYC was detected in HopZ5-YFP and AvrBsT-YFP immuno-precipitates, indicating that HopZ5 and AvrBsT may directly or indirectly associate with JIM2 to activate NbZAR1-dependent defense signaling.

**Fig. 4.**
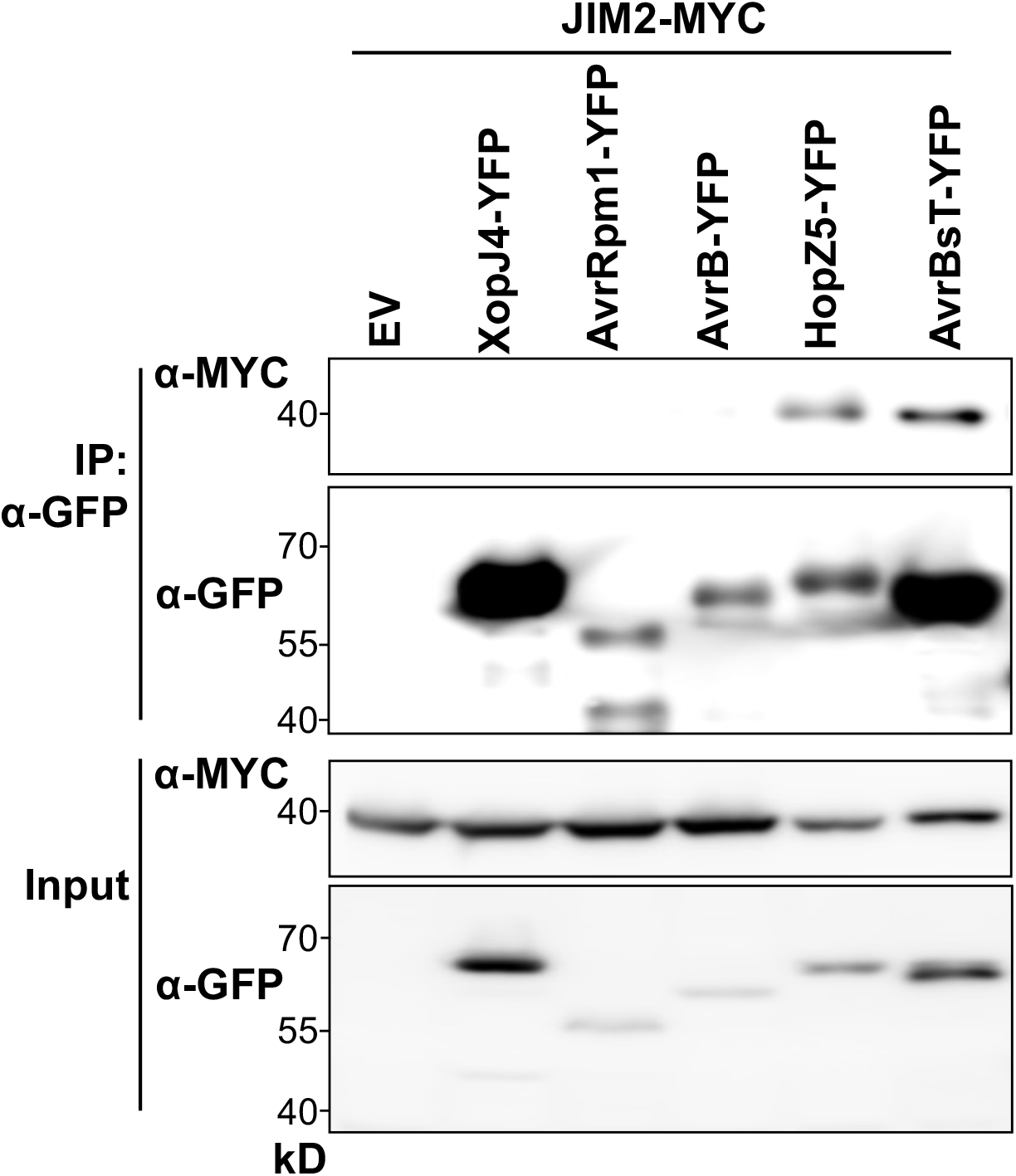
JIM2 co-immunoprecipitates with HopZ5 and AvrBsT. MYC-JIM2 was co-expressed with empty vector (EV), YFP-XopJ4, YFP-AvrRpm1, YFP-AvrB, YFP-HopZ5, or YFP-AvrBsT using *Agrobacterium*. At 48 dpi, proteins were sampled and extracted for co-immunoprecipitation (co-IP) with ⍰-GFP agarose (IP: ⍰-GFP) and immunoblotted with ⍰-MYC and ⍰-GFP. Inputs were collected from the extracted proteins before co-IP.

### Ptr1-dependent recognition of AvrBsT confers bacterial spot disease resistance

The *Xanthomonas* avirulence effector XopQ triggers Roq1-dependent immunity in *N. benthamiana* (Schultink *et al*., 2017). To test if NbPtr1 confers AvrBsT-triggered disease resistance in *N. benthamiana*, we used *Xanthomonas perforans* 4B lacking XopQ (Δ*xopQ*) (Schwartz *et al*., 2015), *X. perforans* Δ*xopQ* lacking *avrBsT* (Δ*xopQ* Δ*avrBsT*) (Schwartz *et al*., 2015), and its complemented strain (Δ*xopQ* Δ*avrBsT* (*avrBsT*)) (for details, see Methods S3) for bacterial growth assay (Fig. 5a). Wild-type and *ptr1* mutant *N. benthamiana* (Nb-1) (for details, see Methods S4) used in this experiment are naturally deficient for ZAR1-mediated recognition of XopJ4 and AvrBsT, possibly due to a lack of JIM2 expression (Fig. S1) (Schultink *et al*., 2019), thus we were able to test NbPtr1-dependent and NbZAR1-independent recognition of AvrBsT. *X. perforans* Δ*xopQ* and Δ*xopQ* Δ*avrBsT* (*avrBsT*) strains showed 100-fold increased growth in *ptr1* mutant compared to wild-type Nb-1 (Fig. 5a). In contrast, the growth of *X. perforans* Δ*xopQ* Δ*avrBsT* strain did not differ in wild-type and *ptr1* mutant *N. benthamiana* (Fig. 5a). These results show that AvrBsT-triggered NbPtr1-mediated immunity effectively restricts *Xanthomonas* growth in *N. benthamiana*.

**Fig. 5.**
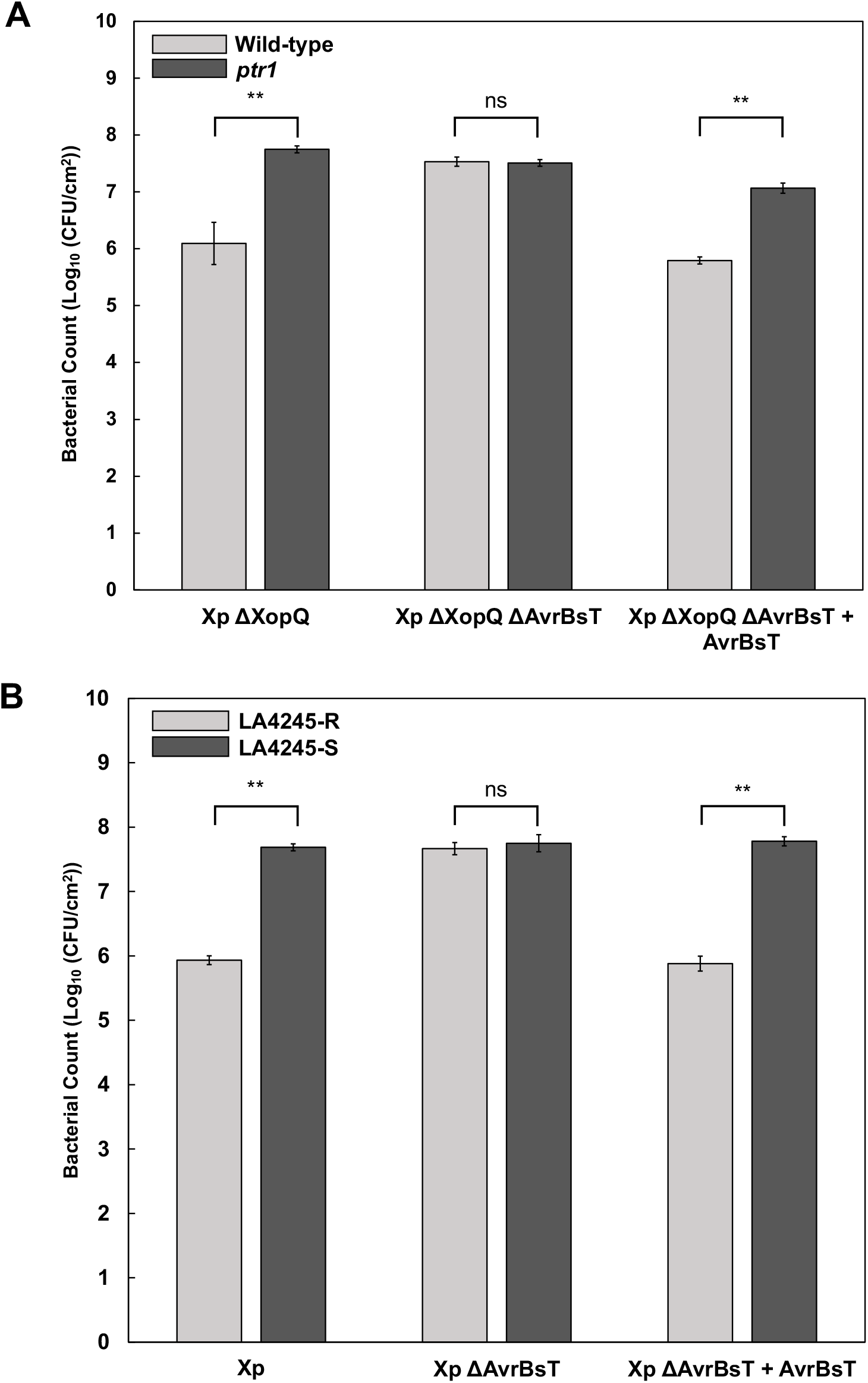
Recognition of AvrBsT by Ptr1 confers bacterial spot disease resistance. (a) AvrBsT restricts *Xanthomonas perforans* growth in wild-type *N. benthamiana* (Nb-1, lacking XopJ4 recognition) but not in *ptr1* knockout mutants. *X. perforans* strains with the indicated genotypes were infiltrated in wild-type or *ptr1 N. benthamiana* at OD_600_ of 0.0001. The growth was assayed at 6 dpi. Error bars indicate ± SD from three biological replicates. Asterisks indicate significant difference, estimated by student’s t-test, between bacterial count in wild-type and *ptr1* N. benthamiana; *(*P* < 0.05) and **(*P* < 0.01). ‘ns’ indicates that there is no significant difference (*P* > 0.05). (b) *X. perforans* strains with the indicated genotypes were infiltrated in tomato introgression lines LA4245-R and LA-4245-S at an OD_600_ of 0.0001. The growth was assayed at 6 dpi. Error bars indicate ± SD from three biological replicates. Asterisks indicate a significant difference, estimated by the student’s t-test, between bacterial count in LA4245-R and LA4245-S; *(*P* < 0.05) and **(*P* < 0.01). ‘ns’ indicates that there is no significant difference (*P* > 0.05).

To test AvrBsT-triggered bacterial growth restriction in tomato, *in planta* growth of *X. perforans* 4B wild-type, *avrBsT* knock out mutant (Δ*avrBsT*) (Schwartz *et al*., 2015), and complemented strain (Δ*avrBsT* (*avrBsT*)) were measured in tomato introgression lines LA4245-R (*Ptr1 ptr1*) and LA4245-S (*ptr1 ptr1*) (Fig. 5b) (Mazo-Molina *et al*., 2020). *X. perforans* wild-type and Δ*avrBsT* (*avrBsT*) showed a 100-fold increase in growth in LA4245-S compared to the growth in LA4245-R (Fig. 5b). No significant difference in growth was observed with *X. perforans* Δ*avrBsT* between LA4245-R and LA4245-S (Fig. 5b). Taken together, these results demonstrate that NbPtr1- and Ptr1-dependent recognition of AvrBsT confer bacterial spot disease resistance in *N. benthamiana* and tomato, respectively.

### Ptr1 and ZAR1 homologs recognize AvrBsT in pepper

AvrBsT triggers immune responses in pepper (Minsavage, 1990; Kim *et al*., 2010). The amino acid sequence identities of Ptr1 and ZAR1 between *N. benthamiana* and pepper homologs are 86.9% and 88.8%, respectively (Fig. S7). Based on our results in *N. benthamiana* and tomato (Fig. 5), we hypothesized that the pepper homologs Ptr1 (CaPtr1) and ZAR1 (CaZAR1) may recognize AvrBsT in pepper. To investigate this, we silenced *CaPtr1* or/and *CaZAR1* gene in pepper using VIGS (Fig. 6a). We used two different *X. campestris* pv. *vesicatoria* strains, Bv5-4a that carries *avrBsT* and Ds1.1 that naturally lacks *avrBsT* to test PCD in pepper (Kim *et al*., 2010). As expected, *X. campestris* pv. *vesicatoria* Ds1.1 did not trigger PCD in all pepper genotypes (Fig. 6a). In contrast, *X. campestris* pv. *vesicatoria* Bv5-4a caused PCD in wild-type, EV-VIGS, CaPtr1-VIGS, and CaZAR1-VIGS peppers (Fig. 6a). Consistent with our *N. benthamiana* results (Fig. 3a), AvrBsT-triggered PCD was abolished in CaPtr1-CaZAR1-VIGS pepper plants (Fig. 6a).

**Fig. 6.**
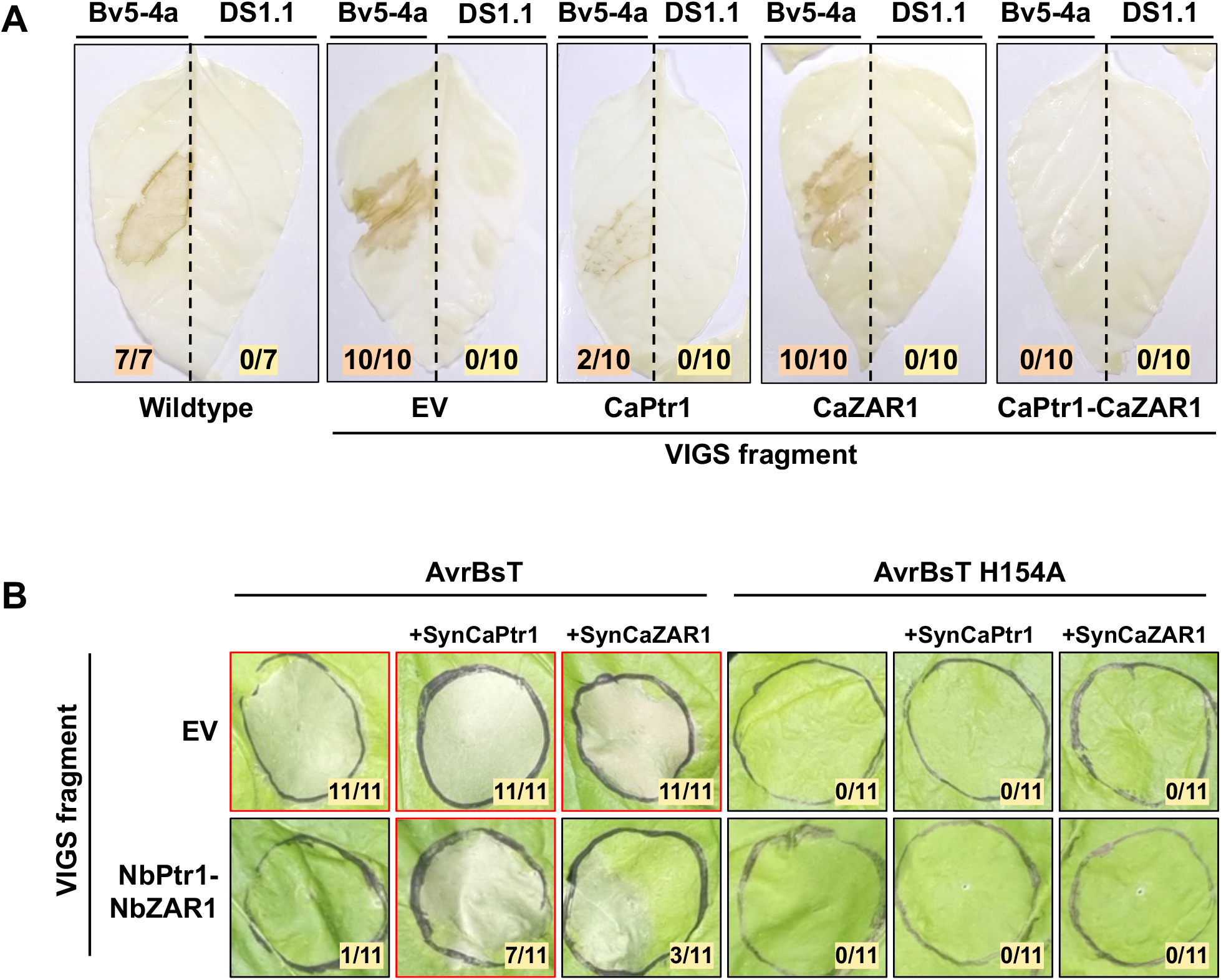
CaPtr1 and CaZAR1 recognize AvrBsT in Pepper. (a) PCD induced by *X. campestris* pv. *vesicatoria* is reduced in CaPtr1-CaZAR1-VIGS peppers. The virulent strain *Xcv* Ds1.1 (lacking AvrBsT) and the avirulent strain Bv5-4a (containing AvrBsT) were infiltrated in wild-type and VIGSed peppers at an OD_600_ of 0.2. The numerator indicates the number of spots with cell death, and the denominator indicates the total number of infiltrations. Photographs of representative methanol-cleared leaves were taken at 3 dpi. (b) Coexpression of SynCaPtr1 but not SynCaZAR1 restores AvrBsT-induced PCD in NbPtr1-NbZAR1-VIGS *N. benthamiana*. AvrBsT (OD_600_ = 0.4), AvrBsT H154A (OD_600_ = 0.4), CaPtr1 (OD_600_ = 0.1), and CaZAR1 (OD_600_ = 0.4) were transiently expressed in *N. benthamiana*. The numerator indicates the number of spots with cell death, and the denominator indicates the total number of infiltrations. If PCD count is higher than 50% of total infiltrations, red borders were marked around the photograph, and black borders were used for lower PCD counts. Photographs of representative leaves were taken at 2 dpi.

To confirm that AvrBsT is recognized by CaPtr1 and CaZAR1 independently, we tested if a transient expression of CaPtr1 or CaZAR1 could restore AvrBsT-induced PCD in NbPtr1-NbZAR1-VIGS *N. benthamiana* (Fig. 6b). Since *CaPtr1* and *CaZAR1* have 89.3% and 90.1% nucleotide identities, respectively, with *N. benthamiana* homologs, codon altered *SynCaPtr1* and *SynCaZAR1* binary constructs were used to evade silencing in NbPtr1-NbZAR1-VIGS plants (Fig. S8, S9). In EV-VIGS plants, agroinfiltration of AvrBsT induced PCD whether SynCaPtr1 or SynCaZAR1 was coexpressed or not (Fig. 6b). In NbPtr1-NbZAR1-VIGS plants, agroinfiltration of AvrBsT alone did not induce PCD, but the coexpression with SynCaPtr1 restored PCD (Fig. 6b). The coexpression of AvrBsT with SynCaZAR1 only weakly restored PCD in NbPtr1-NbZAR1-VIGS plants (Fig. 6b). The expression of the catalytically inactive variant AvrBsT H154A (Cheong *et al*., 2014) did not induce PCD in EV-VIGS and NbPtr1-NbZAR1-VIGS plants suggesting that CaPtr1 or CaZAR1 recognizes AvrBsT acetyltransferase activity (Fig. 6b). Taken together, our *N. benthamiana* and pepper results are consistent and indicate that CaPtr1 plays a major role in the recognition of AvrBsT.

## DISCUSSION

We demonstrate here that NbPtr1 recognizes multiple sequence-unrelated bacterial effectors using the NbNLR VIGS library screening. Furthermore, we reveal that recognition of two YopJ family acetyltransferase effectors, HopZ5 and AvrBsT, is conferred independently by NbZAR1. We also show that JIM2 physically associates with HopZ5 or AvrBsT and might work as a guardee or decoy in NbZAR1-mediated immunity in *N. benthamiana*. Finally, we show that Ptr1 confers bacterial spot disease resistance by recognizing *Xanthomonas*-delivered AvrBsT in *N. benthamiana, S. lycopersicum*, and *C. annum*.

Disease resistance genes are often identified by forward genetic approaches in which a resistant plant is crossed to a susceptible relative or a mutagen is used to induce genetic variation in a resistant population before screening for susceptible individuals and mapping the gene of interest (Schultink & Steinbrenner, 2021). These methods are time and labor-intensive. As NLRs underly most disease resistance traits, reverse genetic approaches focused on *NLR* genes can accelerate resistant gene identification. The improvement of the plant reference genomes and NLR annotation with methods such as SMRT RenSeq facilitates exploration of NLR diversity in a wide range of plant species (Van De Weyer *et al*., 2019; Seong *et al*., 2020). We designed the NbNLR VIGS library based on an improved NLR annotation to perform a rapid reverse genetic screen for NbNLRs mediating recognition of a diverse set of bacterial effector proteins. This work builds upon previous reverse genetic efforts in *Nicotiana* which identified Roq1, RXEG1, and Rpa1 (Schultink *et al*., 2017; Wang *et al*., 2018; Yoon & Rikkerink, 2020). Previously, a hairpin-RNAi library was constructed based on the 345 NbNLRs annotated in an earlier version of *N. benthamiana* genome assembly (Bombarely *et al*., 2012; Brendolise *et al*., 2017). The hairpin library only partially overlaps with the NLRs from our VIGS library which is based on the refined NLR annotations using SMRT-RenSeq (Seong *et al*., 2020) (Table S2), emphasizing the importance of having a complete and high-quality genome annotation for this approach. It is worth noting that the success of forward or reverse genetic approaches can be limited when two or more NLRs redundantly function, as we observed for ZAR1 and Ptr1 recognition of AvrBsT and HopZ5. One way to overcome this functional redundancy is to use a quantitative PCD-scoring method (e.g. ion leakage assay) to detect a decrease rather than a complete loss of elicitor recognition.

The effectors that are recognized by NbPtr1 have been previously associated with modifications of RIN4 in *Arabidopsis*. AvrB and AvrRpm1 induce phosphorylation and HopZ5 and AvrBsT induce acetylation on Thr166 located in the conserved C-terminal NOI domain of RIN4, and these modifications activate RPM1 in *Arabidopsis* (Chung *et al*., 2011; Liu *et al*., 2011; Redditt *et al*., 2019; Choi *et al*., 2021). Furthermore, AvrRpt2 induces proteolytic cleavage on RIN4 and activates RPS2 in *Arabidopsis* (Axtell & Staskawicz, 2003; Mackey *et al*., 2003; Day *et al*., 2005). In tomato, AvrRpt2 is recognized by Ptr1 (Mazo-Molina *et al*., 2020). Overexpression of Ptr1 leads to PCD in *N. glutinosa* and this autoactivation is suppressed by coexpression of tomato RIN4 homologs, and the suppression is released when AvrRpt2 is additionally expressed (Mazo-Molina *et al*., 2020). Thus, it is plausible that the effectors that are recognized by NbPtr1 may modify *N. benthamiana* RIN4 homologs as their guardees or decoys. However, silencing NbRIN4 homologs with VIGS did not have any effect on effector recognition in *N. benthamiana* (Fig. S6). Therefore, it is unclear that RIN4 is required for effector-triggered NbPtr1 activation. RIN4 has two structured regions characterized as N-terminal and C-terminal NOI domains (Lee *et al*., 2015). These domains are functionally important and conserved in other proteins called NOI proteins (Afzal *et al*., 2011; Toruño *et al*., 2019). NOI proteins besides RIN4 may be involved in NbPtr1 activation since some of the effectors have been shown to act on multiple NOI proteins. For example, AvrRpm1 can ADP-ribosylate RIN4 and ten additional NOI proteins that lack sequence similarity with RIN4 other than the presence of NOI domains in *Arabidopsis* (Redditt et al., 2019). This suggests that NOI proteins can be enzymatically modified by the RIN4-targeting effectors. Taken together, *N. benthamiana* NOI proteins whether redundantly with NbRIN4 homologs may be targeted by the effectors and are monitored by NbPtr1. It would be interesting to investigate if NOI proteins are required for NbPtr1 function in recognizing multiple bacterial effectors in *Solanaceae* species.

HopZ5 and AvrBsT recognition is conferred by overlapping function of Ptr1 and ZAR1 in *N. benthamiana* suggesting that there could be multiple host targets of these effectors (Fig. 7). We reveal that the RLCK XII family protein, JIM2, is required for HopZ5 and AvrBsT-triggered immunity and physically associates with HopZ5 and AvrBsT (Fig. 3 and 4). Although JIM2 is also required for recognition of XopJ4 (Fig. 7), previously performed co-IP (Schultink *et al*., 2019) and our result (Fig. 4) show that JIM2 interaction is significantly weaker with XopJ4 than with HopZ5 or AvrBsT. This suggests that the mechanism by which ZAR1 recognizes XopJ4 may be distinct from that which it uses to recognize HopZ5 and AvrBsT. ZAR1 is known to utilize different RLCK proteins for recognizing diverse effectors in *Arabidopsis*, with all requiring an RLCK XII protein and at least some requiring an additional RLCK VII protein (Lewis *et al*., 2010; Wang *et al*., 2015; Seto *et al*., 2017; Liu *et al*., 2019; Schultink *et al*., 2019; Martel *et al*., 2020). As XopJ4 and AvrBsT show only 38% identity, it would not be surprising if they have different virulence targets and could be recognized by different guardee or decoy proteins. For instance, XopJ4 may directly interact with an unknown RLCK VII to activate ZAR1-mediated immunity with JIM2, whereas JIM2 could be directly acetylated by HopZ5 and AvrBsT to activate ZAR1-mediated immunity.

**Fig. 7.**
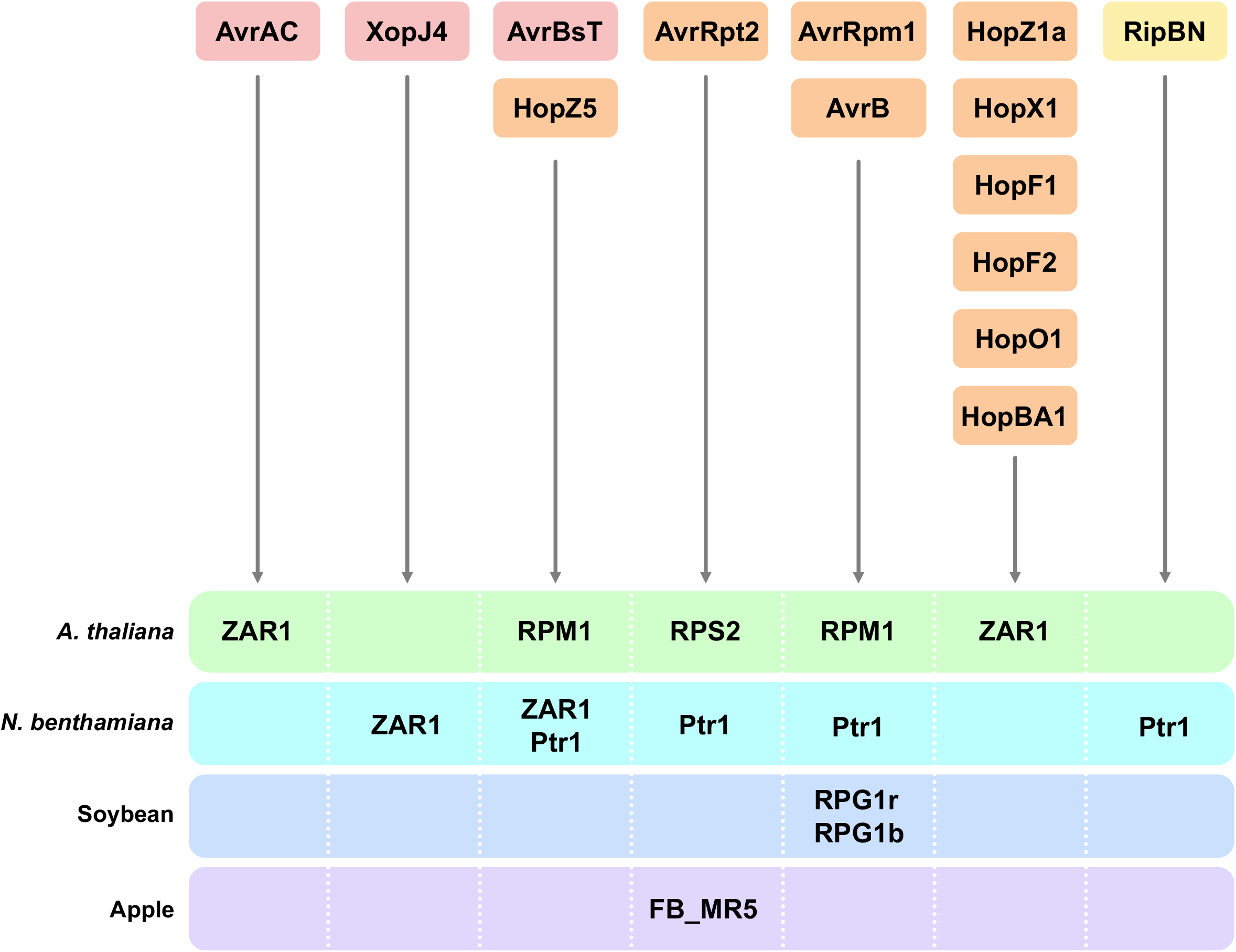
Convergent evolution of effector recognition in diverse plant species. AvrAC from *X. campestris* is recognized by ZAR1 in *A. thaliana* (Wang *et al*., 2015). XopJ4 from *X. perforans* is recognized by ZAR1 in *N. benthamiana* (Schultink et al., 2019). AvrBsT from *X. perforans* and HopZ5 from *P. syringae* pv. *actinidiae* are recognized by RPM1 (Choi *et al*., 2021) in *A. thaliana* and ZAR1 and Ptr1 in *N. benthamiana* (this study) (Mazo-Molina *et al*., 2020). AvrRpt2 from *P. syringae* pv. *tomato* is recognized by RPS2 in *A. thaliana* (Axtell & Staskawicz, 2003; Mackey *et al*., 2003) and Ptr1 in *N. benthamiana* (this study), and its *Erwinia amylovora* homolog by FB_MR5 in apple (Vogt *et al*., 2013; Prokchorchik *et al*., 2020). AvrRpm1 from *P. syringae* pv. *maculicola* and AvrB from *P. glycinea* are recognized by RPM1 in *A. thaliana* (Chung *et al*., 2011; Liu *et al*., 2011; Redditt *et al*., 2019), Ptr1 in *N. benthamiana* (this study), and RPG1r or RPG1b, respectively, in soybean (Ashfield *et al*., 1995; Ashfield *et al*., 2004). HopZ1a, HopX1, HopF1, HopF2, HopO1, and HopBA1 from *Pseudomonas* are recognized by ZAR1 in *A. thaliana* (Lewis *et al*., 2010; Laflamme *et al*., 2020; Martel *et al*., 2020). RipBN from *R. solanacearum* is recognized by Ptr1 (Mazo-Molina et al., 2020). *Xanthomonas* effectors are in red, *Pseudomonas* effectors are in orange, and *Ralstonia* effectors are in yellow.

Maintaining multiple NLRs that overlap in effector recognition is costly for the plant survival (Tian *et al*., 2003), unless these NLRs have been evolutionarily selected to provide resilience in resistance against coevolving pathogens. Sensor NLRs recognize effectors and activate downstream immune responses which are mediated by a network of helper NLRs (Wu *et al*., 2017; Jacob *et al*., 2021; Ngou *et al*., 2022b). In addition, EP (EDS1-PAD4 specific domain) proteins such as EDS1, PAD4, and SAG101 work in coordination with helper NLRs and execute resistance against pathogens (Feehan *et al*., 2020; Huang *et al*., 2022). These helper NLRs and downstream signaling components are attractive virulence targets for pathogen effector proteins as inactivation of them could interfere with sensor NLR function and allow a pathogen to evade effector-triggered immunity. Indeed, effectors from the late blight pathogen, *Phytophthora infestans*, and the cyst nematode, *Globodera rostochiensis*, were shown to target the NRC network to suppress either Prf or Rpi-blb2-mediated immunity (Derevnina *et al*., 2021). Since NLR-mediated immunity can be suppressed by effectors, it would be advantageous for host plants to harbor NLRs that have expanded effector recognition or associate with diverse helper NLRs. Therefore, overlapping effector recognition of Ptr1 and ZAR1 may be a consequence of coevolving bacterial effectors that can suppress NLRs (Fig. 7). Furthermore, enhanced resistance conferred by broad and overlapping effector recognition of Ptr1 and ZAR1 is desirable for crop engineering. *X. perforans* strains that recently caused major outbreaks in tomato fields carry *avrBsT* (Schwartz *et al*., 2015; Timilsina *et al*., 2016), but *S. lycopersicum* lacks functional Ptr1 and ZAR1 (Schultink *et al*., 2019; Mazo-Molina *et al*., 2020). Since *X. perforans* carrying *avrBsT* shows a reduced virulence in tomato introgression line LA4245-R (*Ptr1 ptr1*) (Fig. 5b), expressing functional *Ptr1* and *ZAR1* in *S. lycopersicum* may confer resistance against newly emerging strains of *X. perforans*. These genes could be stacked in tomato along with *EFR* from *Arabidopsis, Bs2* from pepper, and *Roq1* from *Nicotiana* to confer durable resistance against multiple bacterial pathogens (Kunwar *et al*., 2018; Thomas *et al*., 2020). In addition to introducing Ptr1 and ZAR1 to susceptible cultivars, further information about the host targets of the avirulence effectors including JIM2 would be useful for crop engineering.

## ACKNOWLEDGEMENTS

This study was carried out with the support of the National Research Foundation of Korea (NRF) grant (NRF-2018R1A5A1023599 and 2019R1A2C2084705) funded by the Korean government (MSIT) and US National Science Foundation grant IOS-1546625 (GBM). YJA and HK were supported by the BK21 funded by the Ministry of Education, Republic of Korea (4120200313623). We thank Liam Clearly for maintaining the Nb1-*ptr1* seeds.

## AUTHOR CONTRIBUTIONS

YJA, HK, SC, and KHS designed the study. YJA, HK, SC, CMM, MP, BK, HY, NZ, and AS performed the experiments. YJA, HK, SC, CMM, CS, AS, GBM, and KHS analyzed the data. YJA, HK, and KHS wrote the manuscript.

## DATA AVAILABILITY

All relevant data from this study are included within the manuscript and its supporting materials.

## SUPPORTING INFORMATION

**Fig. S1** XopJ4 recognition in Nb-0, Nb-1, and Nb-1 *ptr1* CRISPR mutant.

**Fig. S2** NbPtr1 silencing by multiple VIGS fragments.

**Fig. S3** Ptr1 mediates the recognition of AvrRpt2, AvrRpm1, AvrB, HopZ5, and AvrBsT in *N. glutinosa*.

**Fig. S4** Nucleotide Alignment of *NbPtr1* and *SynPtr1*.

**Fig. S5** Nucleotide alignment of *NbZAR1* and *SynZAR1*.

**Fig. S6** NbRIN4-1,2,3-VIGS does not abolish the effector recognition in *N. benthamiana*.

**Fig. S7** Amino acid alignment of NbPtr1 and NbZAR1 with their pepper homologs.

**Fig. S8** Nucleotide alignment of *CaPtr1* and *SynCaPtr1*.

**Fig. S9** Nucleotide alignment of *CaZAR1* and *SynCaZAR1*.

**Table S1** Sequence and NLR information of the NbNLR VIGS library.

**Table S2** Comparisons of NLRs between the NbNLR VIGS library and the hairpin library from Brendolise *et al*., 2017.

**Table S3** Primers used in this study.

**Methods S1** RNA extraction and quantitative RT-PCR

**Methods S2** Deletion of *hopQ1-1* from *Pseudomonas syringae* pv. *tomato* DC3000

**Methods S3** AvrBsT complementation of *Xanthomonas perforans* 4B _Δ_*xopQ* _Δ_*avrBsT*

**Methods S4** Generation of *Nicotiana benthamiana ptr1* mutant using CRISPR/Cas9

**Methods S5** *P. syringae* pv. *tomato* Δ*hopQ1-1* Δ*avrPto* Δ*avrPtoB* culture and transformation

**Methods S6** *P. syringae* pv. *tomato* inoculation and population assays in tomato

**Supporting Reference**

## Notes

### Competing Interest Statement

The authors have declared no competing interest.

