## Supplementary Figures and Methods for "Ptr1 and ZAR1 immune receptors confer overlapping and distinct bacterial pathogen effector specificities"

The following Supporting Information is available for this article:

**Fig. S1** XopJ4 recognition in Nb-0, Nb-1, and Nb-1 *ptr1* CRISPR mutant.

**Fig. S2** NbPtr1 silencing by multiple VIGS fragments.

**Fig. S3** Ptr1 mediates the recognition of AvrRpt2, AvrRpm1, AvrB, HopZ5, and AvrBsT in *N. glutinosa*.

**Fig. S4** Nucleotide Alignment of *NbPtr1* and *SynPtr1*.

**Fig. S5** Nucleotide alignment of *NbZAR1* and *SynZAR1*.

**Fig. S6** NbRIN4-1,2,3-VIGS does not abolish the effector recognition in *N. benthamiana*.

**Fig. S7** Amino acid alignment of NbPtr1 and NbZAR1 with their pepper homologs.

**Fig. S8** Nucleotide alignment of *CaPtr1* and *SynCaPtr1*.

**Fig. S9** Nucleotide alignment of *CaZAR1* and *SynCaZAR1*.

**Table S1** Sequence and NLR information of the NbNLR VIGS library.

**Table S2** Comparisons of NLRs between the NbNLR VIGS library and the hairpin library from Brendolise *et al*., 2017.

**Table S3** Primers used in this study.

**Methods S1** RNA extraction and quantitative RT-PCR

**Methods S2** Deletion of *hopQ1-1* from *Pseudomonas syringae* pv. *tomato* DC3000

**Methods S3** AvrBsT complementation of *Xanthomonas perforans* 4B *ΔxopQ ΔavrBsT*

**Methods S4** Generation of *Nicotiana benthamiana ptr1* mutant using CRISPR/Cas9

**Methods S5** *P. syringae* pv. *tomato* Δ*hopQ1-1* Δ*avrPto* Δ*avrPtoB* culture and transformation

**Methods S6** *P. syringae* pv. *tomato* inoculation and population assays in tomato

**Supporting Reference**

**Fig. S1** XopJ4 recognition in Nb-0, Nb-1, and Nb-1 *ptr1* CRISPR mutant. (a) *N. benthamiana* were agroinfiltrated with AvrRpt2, AvrRpt2 C122A, HopZ1a, ZED1, XopJ4, AvrBsT, and HopZ5 with an $OD_{600}$ of 0.4. Photographed at 4 dpi. (b) *N. benthamiana* were agroinfiltrated with GFP, XopJ4, ZAR1, and JIM2 with an $OD_{600}$ of 0.5. Photographed at 4 dpi. (c) JIM2 expression in wild-type Nb-0 and Nb-1 was measured using semi-quantitative PCR. *NbActin* expression was used as a reference (24 cycles). *JIM2* was amplified with 32 cycles.

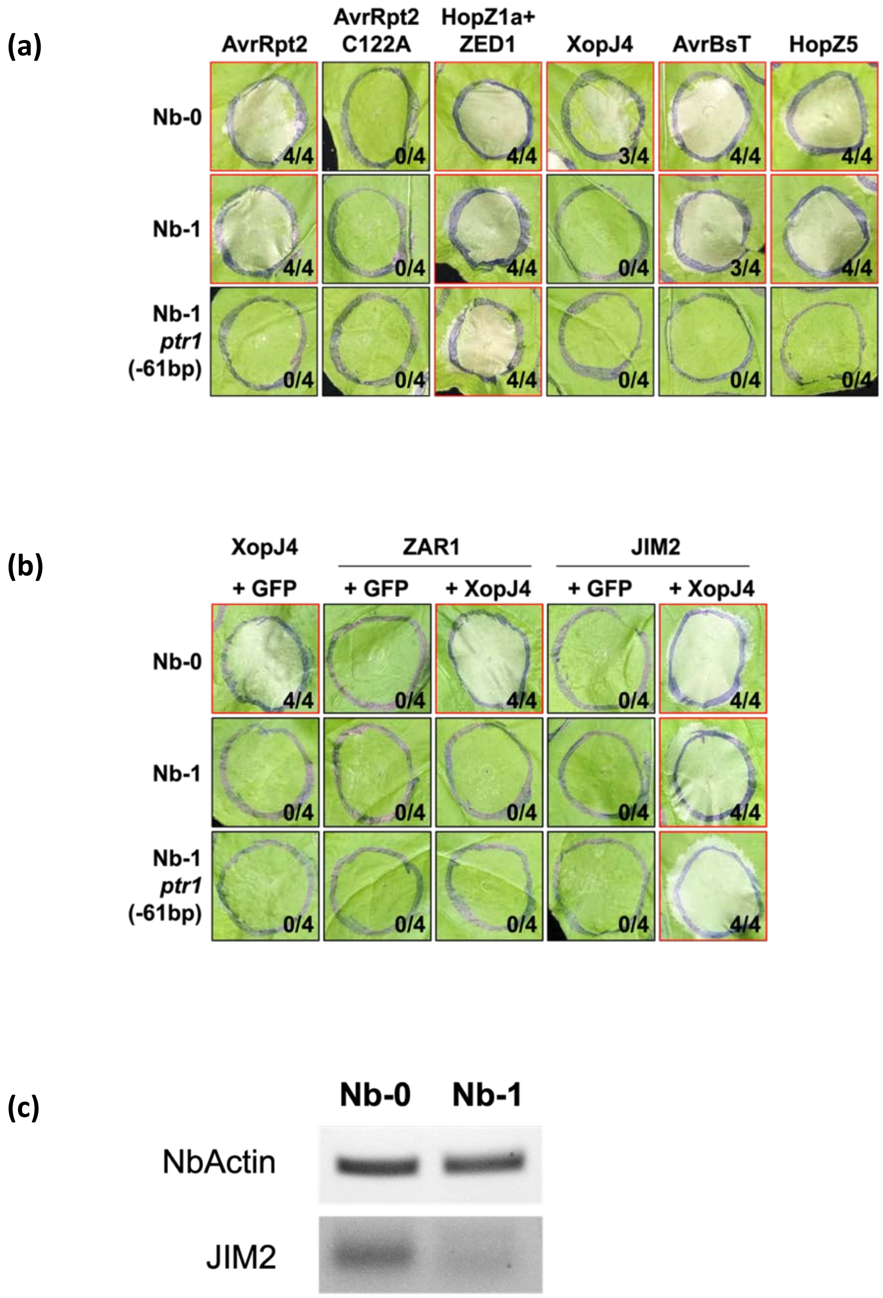

**Fig. S2** NbPtr1 silencing by multiple VIGS fragments. CDS of NbPtr1 (Niben101Scf07061g00004.1) and its close homolog in *N. benthamiana* (Niben101Scf00597g00011.1) are aligned, and SNPs between the two genes are highlighted in black. NbPtr1 and its homolog have 88.9% nucleotide identity. Brown annotations indicate VIGS fragments designed for NbPtr1 silencing. Com-49-3 VIGS targets from 71 to 190 nucleotides in the *NbPtr1* gene, and NbPtr1-VIGS targets the first 300 nucleotides of *NbPtr1*. Note that Com-49-3 VIGS is partially located in the intron of NbPtr1 homolog.

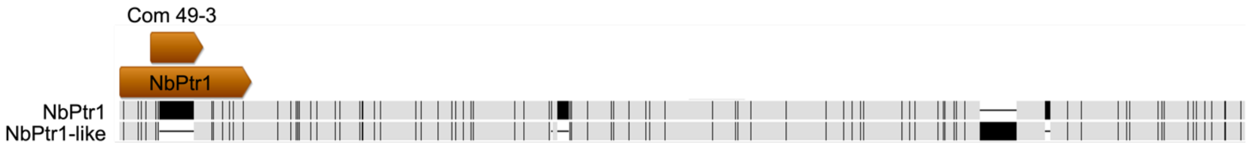

**Fig. S3** Ptr1 mediates the recognition of AvrRpt2, AvrRpm1, AvrB, HopZ5, and AvrBsT in *N. glutinosa*. *N. glutinosa* leaves were syringe-infiltrated with *Agrobacterium* strains carrying *Ptr1:HA* ($OD_{600}$ = 0.025), and *AvrBsT:HA*, *AvrBsT H154A:FLAG*, *AvrRpm1:FLAG*, *AvrB:HA*, *HopZ5:HA* or *AvrRpt2:Myc* ($OD_{600}$ = 0.05). Photographs were taken 48 hr after infiltration and are representative of two independent experiments.

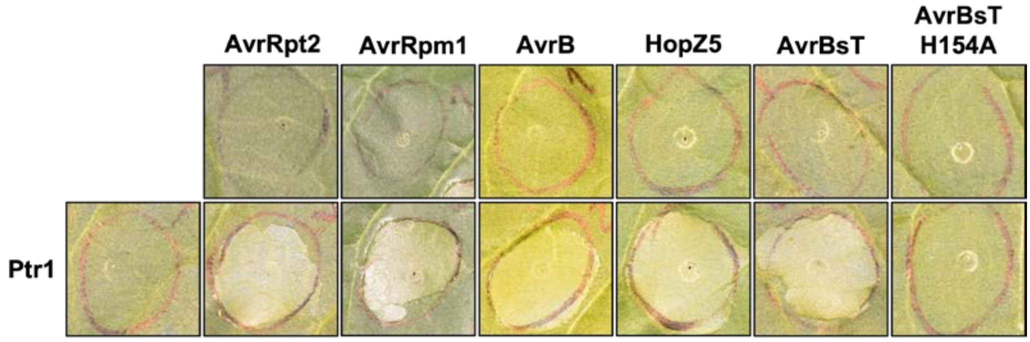

**Fig. S4** Nucleotide alignment of *NbPtr1* and *SynPtr1*. Identical nucleotides are highlighted in black. NbPtr1 and SynPtr1 have the same amino acid sequence.

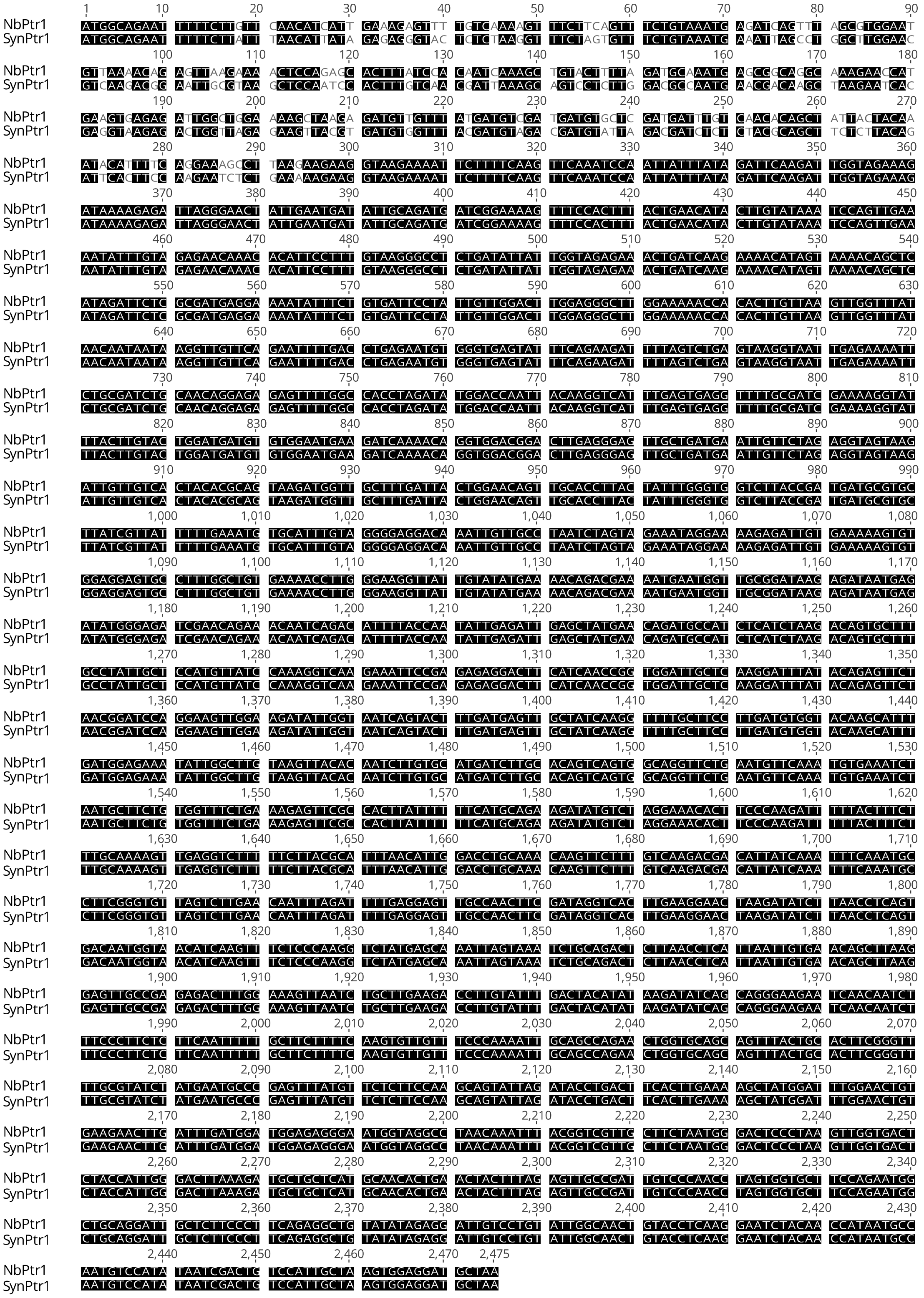

**Fig. S5** Nucleotide alignment of *NbZAR1* and *SynZAR1*. Identical nucleotides are highlighted in black. NbZAR1 and SynZAR1 have the same amino acid sequence.

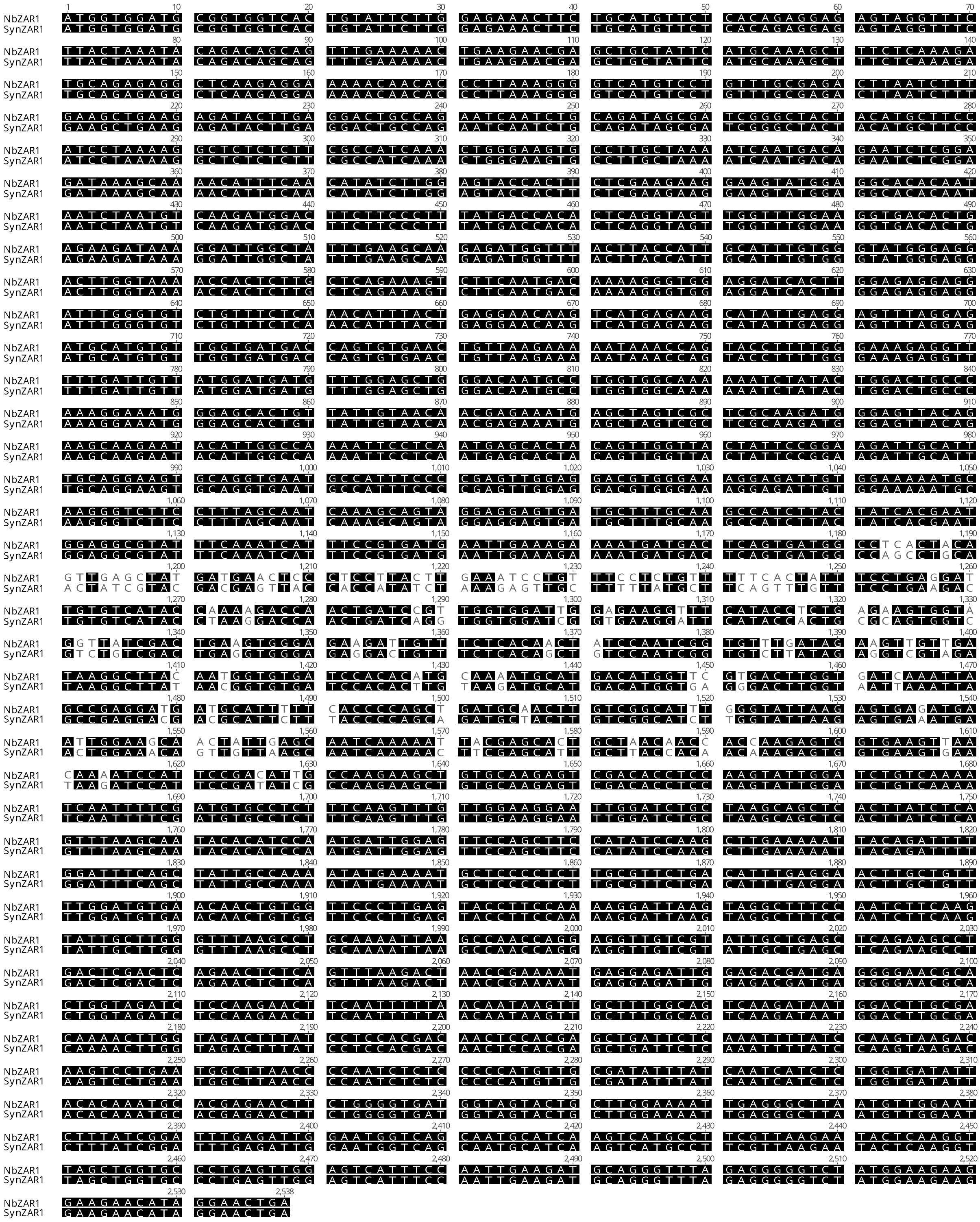

**Fig. S6** NbRIN4-1,2,3-VIGS does not abolish effector recognition in *N. benthamiana*. (a) EV-VIGS, NbPtr1-VIGS, and NbRIN4-1,2,3-VIGS *N. benthamiana* were agroinfiltrated with AvrRpt2, AvrRpt2 C122A, AvrB, AvrRpm1, HopZ5, and AvrBsT with an $OD_{600}$ of 0.4. Red and black borders indicate the presence or the absence of PCD, respectively. The numerator indicates the number of spots with cell death, and the denominator indicates the total number of infiltrations. Photographed at 2 dpi. Similar results were shown at least three times. (b) Expression of NbRIN4 homologs in wild-type, EV-VIGS, NbPtr1-VIGS, NbRIN4-1,2,3-VIGS plants was measured using semi-quantitative PCR. *NbActin* expression was used as a reference (27 cycles). *TRV CP* was amplified with 23 cycles. *NbPtr1* was amplified with 32 cycles. *NbRIN4-1*, *NbRIN4-2*, and *NbRIN4-3* were amplified with 27 cycles. Accession numbers for NbRIN4 genes are NbRIN4-1 (Genbank: KX272617.1), NbRIN4-2 (Solgenomics: Niben101Scf08799g00001), and NbRIN4-3 (Solgenomics: Niben101Scf03488g06005.1) (Prokchorchik *et al.*, 2020).

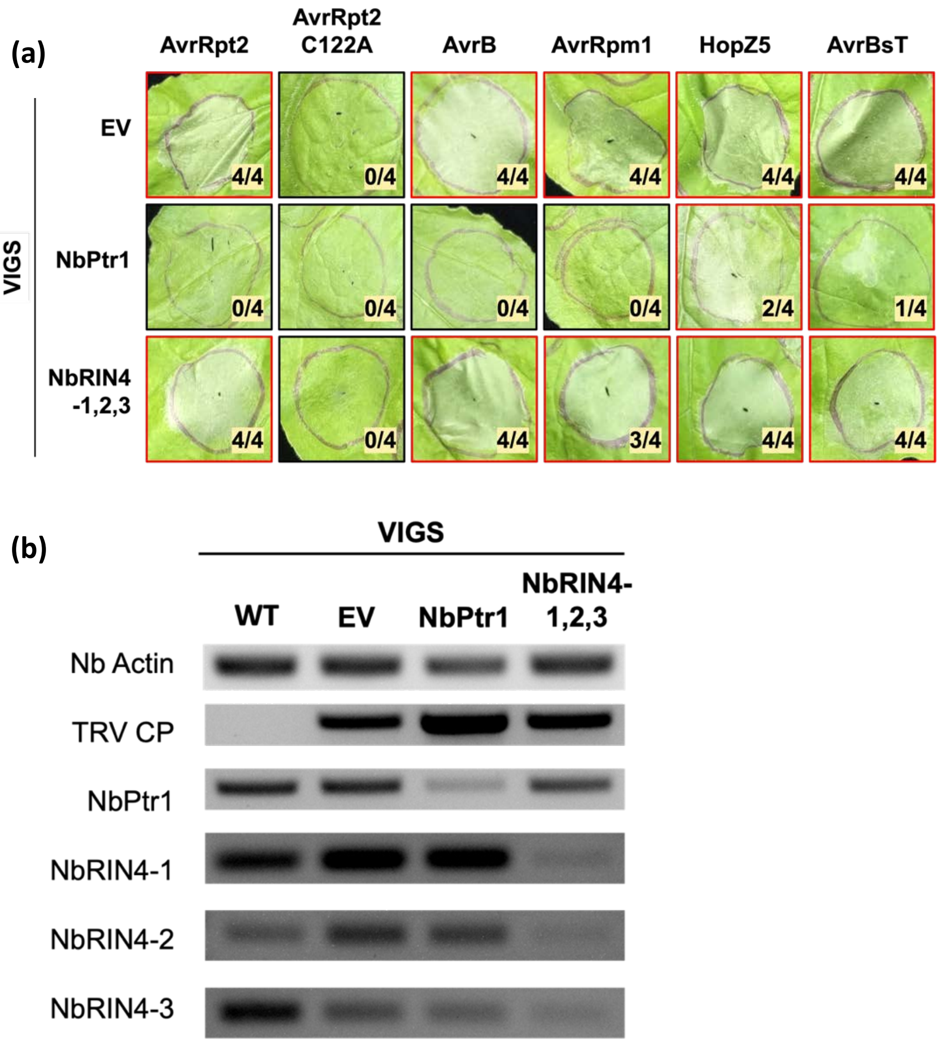

**Fig. S7** Amino acid alignment of NbPtr1 and NbZAR1 with their pepper homologs.

**(a)** NbPtr1 and CaPtr1 (Solgenomics ID: Ca05g0030) (Mazo-Molina *et al.*, 2020) have 86.9% amino acid identity. Identical amino acids shown in black.

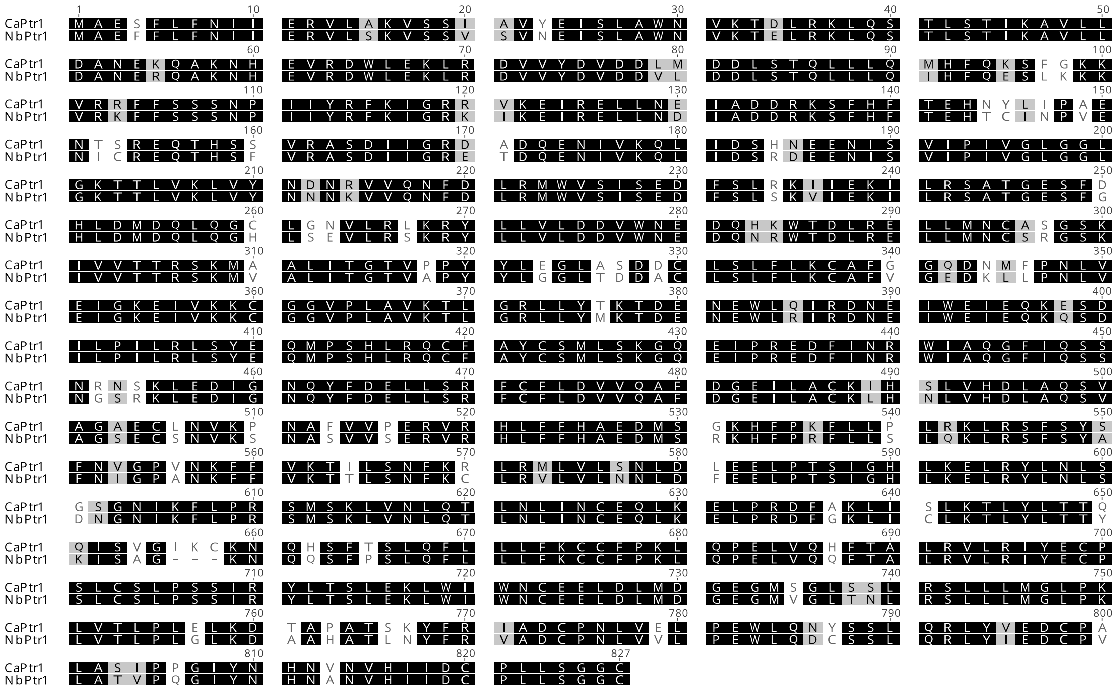

**(b)** NbZAR1 and CaZAR1 (Genbank: XM_016705191) (Schultink *et al.*, 2019) have 88.8% amino acid identity. Identical amino acids shown in black.

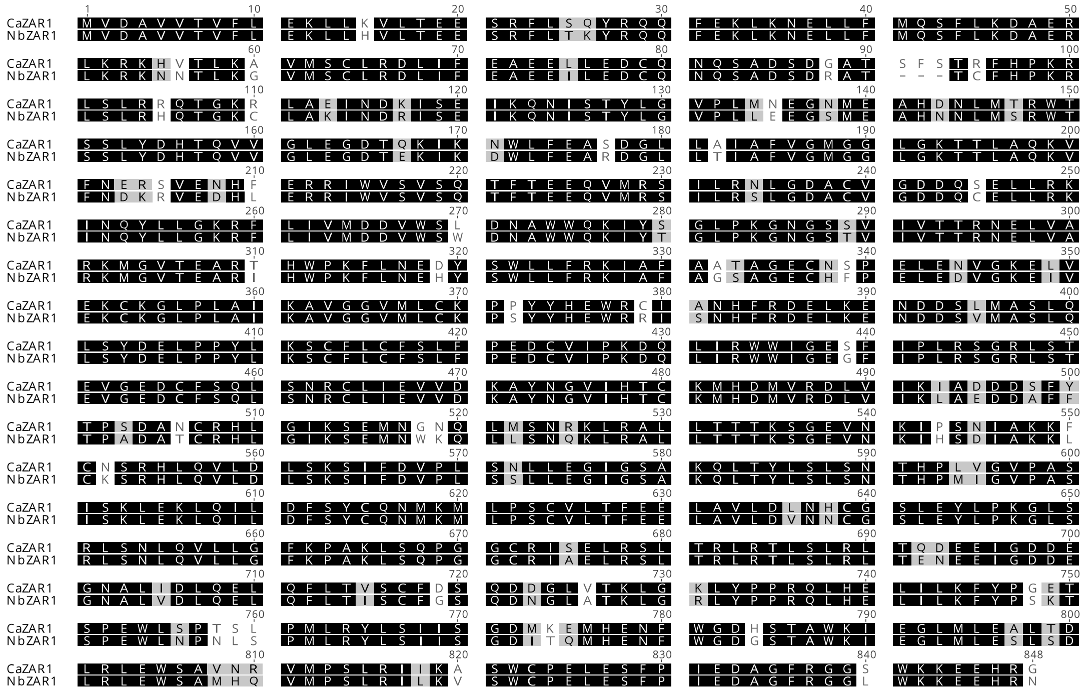

**Fig. S8** Nucleotide alignment of *CaPtr1* and SynCaPtr1. Identical nucleotides are highlighted in black. Both have the same amino acid sequence.

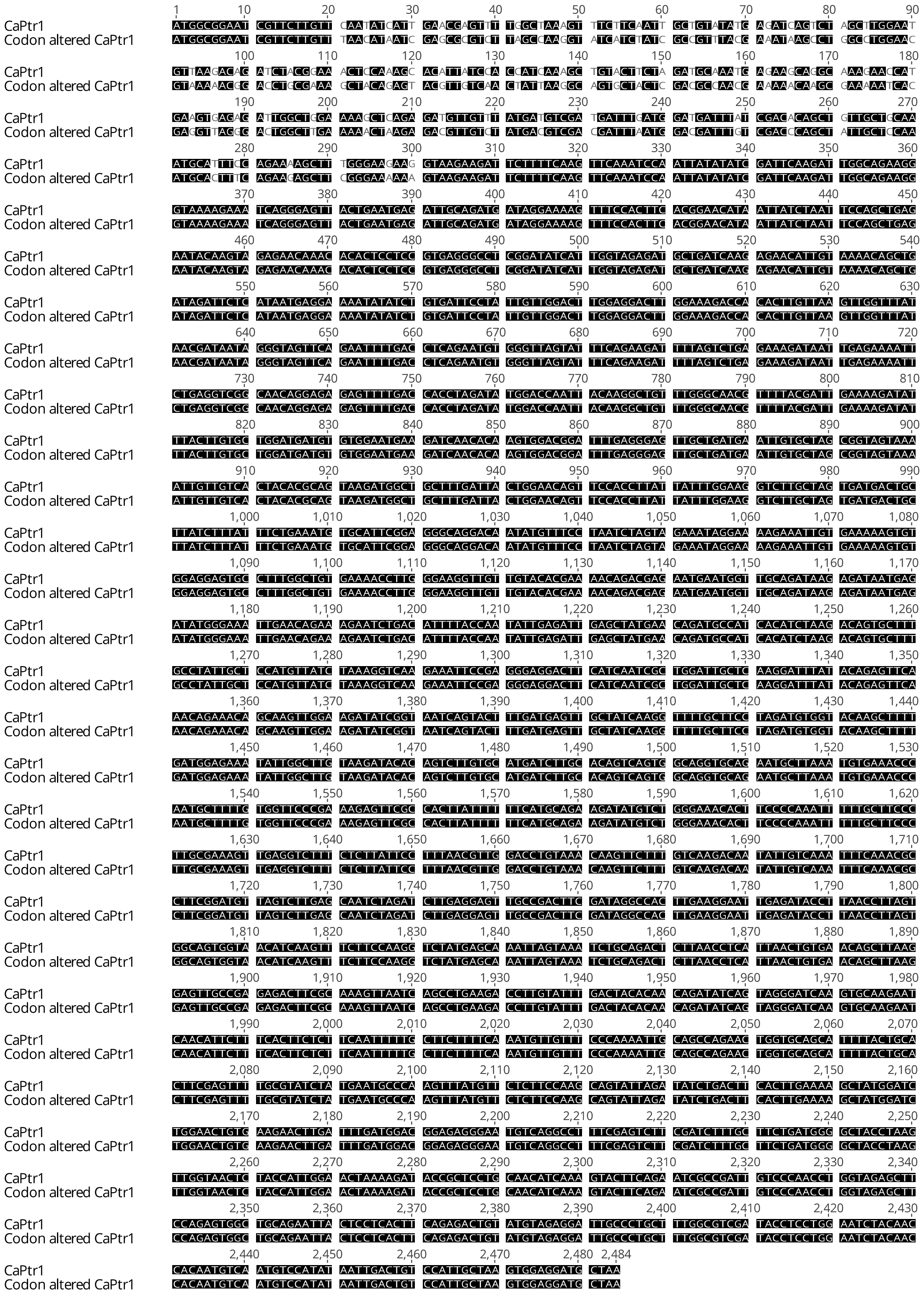

**Fig. S9** Nucleotide alignment of *CaZAR1* and *SynCaZAR1*. Identical nucleotides are highlighted in black. Both have the same amino acid sequence.

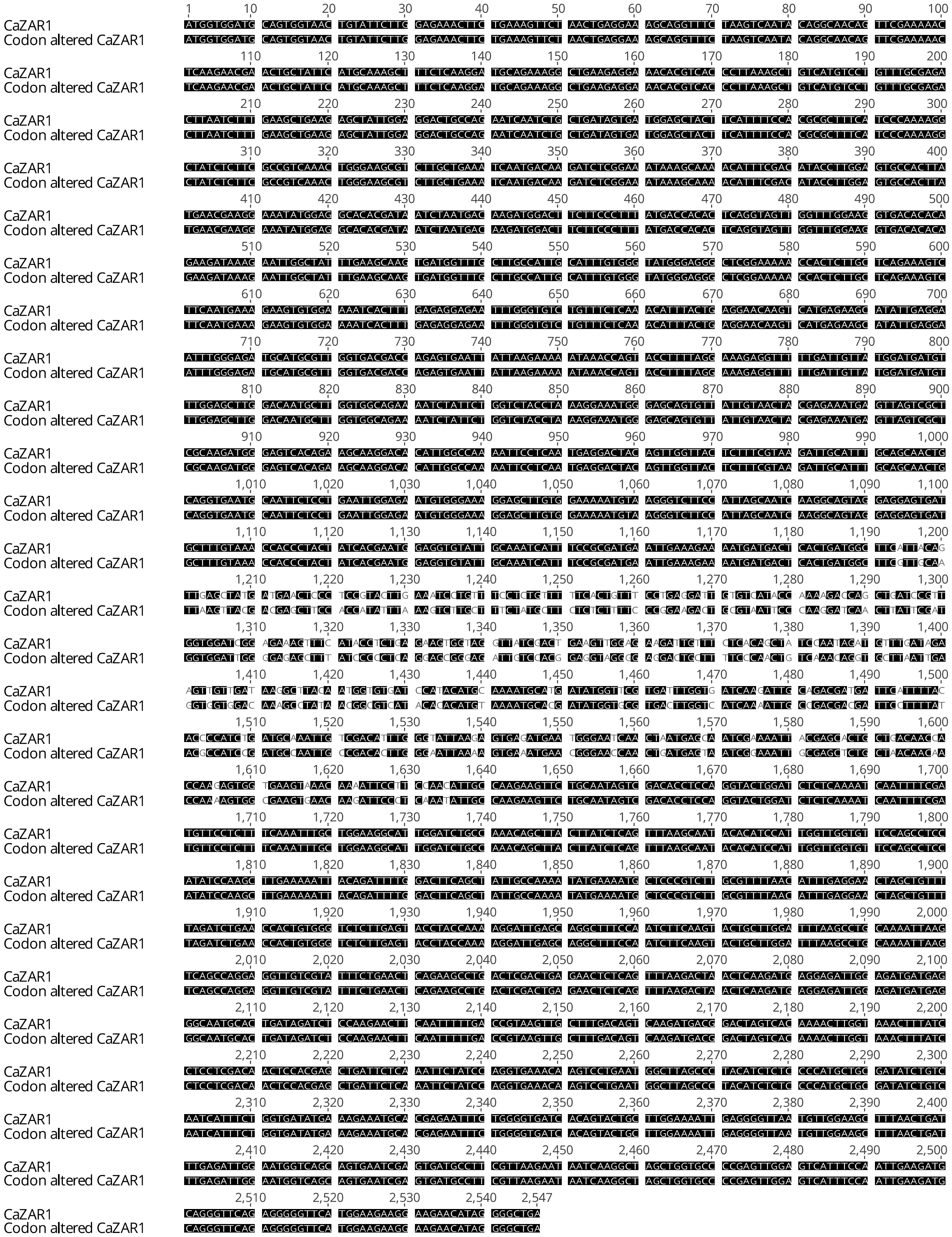

**Table S3** Primers used in this study.

| Gene | Identifier | Forward / Reverse | Sequence | Source |
|---|---|---|---|---|
| **Primers used for genotyping** | | | | |
| *NbPtr1* | PKSP 5499 | Forward | GTGCTCGATGATTTGTCAAC | This work |
| *NbPtr1* | PKSP 5500 | Reverse | TCTTCTGAAATACTCACCCAC | This work |
| Introgressed region in Chr4 69.13 Mb | Spenn-ch04_5416619 | Forward | TAATGAGGCAGAGCAAGTTT | Mazo-Molina et al. (2019) |
| Introgressed region in Chr4 69.13 Mb | Spenn-ch04_5416620 | Reverse | CCCTCAAGAACCATGAATCA | Mazo-Molina et al. (2019) |
| *hopQ1-1* | PKSP 5621 | Forward | ATCGATATCATGCAGCGCTTCAA | This work |
| *hopQ1-1* | PKSP 5622 | Reverse | CGTTTCGGTTTCATCACCAGGG | This work |
| *hopQ1-1* | PKSP 5623 | Reverse | GAAGCTTGGATATGGTGAGGCTC | This work |
| *xopJ4* | oCM296 | Forward | ATGAAAAACATATTTAGGTCACTTGGTCTC | This work |
| *xopJ4* | oCM297 | Reverse | TTAGCTACGACTCAACGCATGAC | This work |
| **Primers used for cloning modules for Golden Gate assembly** | | | | |
| *xopJ4* | PKSP 4719 | Forward | GGTCTCAAATGAAAAACATATTTAGGTCACTTGGGC | This work |
| *xopJ4* | PKSP 5346 | Reverse | GGTCTCTCGAAACTAGTGCTACGACTCAACG | This work |
| *avrRpm1* | PKSP 4736 | Forward | GGTCTCAAATGGGCTGTGTATCGAGCA | This work |
| *avrRpm1* | PKSP 4737 | Reverse | GGTCTCACGAAAAAGTCATCTTCTGAGTCAGACTGA | This work |
| *SynPtr1* | PKSP 3937 | Forward | GGTCTCTAATGGCAGAATTTTTCTTGTTCAACA | This work |
| *SynPtr1* | PKSP 3938 | Reverse | GGTCTCTACAAAGGAATGTGTTTGTTCTCTAC | This work |
| *SynPtr1* | PKSP 3939 | Forward | GGTCTCATTGTAAGGGCCTCTGATATTATTGG | This work |
| *SynPtr1* | PKSP 3940 | Reverse | GGTCTCTCCTCAACTTTTGCAAAGAAAGTAAAAA | This work |
| *SynPtr1* | PKSP 3941 | Forward | GGTCTCAGAGGTCTTTTTCTTACGCATTTAACA | This work |
| *SynPtr1* | PKSP 3942 | Reverse | GGTCTCTCGAAGCATCCTCCACTTAGCAATGG | This work |
| *SynZAR1* | PKSP 4485 | Forward | GGTCTCGAATGGTGGATGCGGTGGT | This work |
| *SynZAR1* | PKSP 4486 | Reverse | GGTCTCACCATCTTGCGAGCGACTAG | This work |
| *SynZAR1* | PKSP 4487 | Forward | GGTCTCTATGGGAGTTACAGAAGCAAGAATAC | This work |
| *SynZAR1* | PKSP 4488 | Reverse | GGTCTCACTTGGATATGGAAGCTGGAACTC | This work |
| *SynZAR1* | PKSP 4489 | Forward | GGTCTCGCAAGCTTGAAAAATTACAGATTTTGGATT | This work |
| *SynZAR1* | PKSP 4490 | Reverse | GGTCTCACGAAGTTCCTATGTTCTTCCTTCTTCCATA | This work |

| | | | | |
|---|---|---|---|---|
| *CaPtr1* | PKSP 5210 | Forward | GGTCTCAAATGGCGGAATCGTTCTTG | This work |
| *CaPtr1* | PKSP 5211 | Reverse | GGTCTCTACCTTCCAAATAATAAGGTGGAAC | This work |
| *CaPtr1* | PKSP 5212 | Forward | GGTCTCAAGGTCTTGCTAGTGATGACTGCTTAT | This work |
| *CaPtr1* | PKSP 5213 | Reverse | GGTCTCTAAGTGGCCTATCGAAGTCGGC | This work |
| *CaPtr1* | PKSP 5214 | Forward | GGTCTCAACTTGAAGGAATTGAGATACCTTAACCT | This work |
| *CaPtr1* | PKSP 5215 | Reverse | GGTCTCTCGAAGCATCCTCCACTTAGCAA | This work |
| *CaZAR1* | PKSP 5216 | Forward | GGTCTCAAATGGTGGATGCAGTGGTAA | This work |
| *CaZAR1* | PKSP 5217 | Reverse | GGTCTCTTTGAGGAATTTTGGCCAATGT | This work |
| *CaZAR1* | PKSP 5218 | Forward | GGTCTCATCAATGAGGACTACAGTTGGTTACTC | This work |
| *CaZAR1* | PKSP 5219 | Reverse | GGTCTCTGAGATCCACAGTGGTTCAGATCTAAAA | This work |
| *CaZAR1* | PKSP 5220 | Forward | GGTCTCATCTCTTGAGTACCTACCAAAAGGATTG | This work |
| *CaZAR1* | PKSP 5221 | Reverse | GGTCTCTCGAAGCCCCTATGTTCTTCCTTC | This work |
| *JIM2* | PKSP 4956 | Forward | GGTCTCAAATGGATTGCATAAAGAAGATGTGG | This work |
| *JIM2* | PKSP 4957 | Reverse | GGTCTCTCGAACTCAAATTGCAGGATCTTTCTTG | This work |
| **Primers used for cloning modules for Golden Gate assembly into pTRV2-66** | | | | |
| Com49-1 | PKSP 4627 | Forward | GGTCTCAAGGTATGAAGTCAAAGTTGCAAAAGCT | This work |
| Com49-1 | PKSP 4628 | Reverse | GGTCTCTAAGCGAAAGAACCTTTTTTTAGAGACACATTC | This work |
| Com49-2 | PKSP 4629 | Forward | GGTCTCAAGGTTATCGAACAAGAAGCACCAGATT | This work |
| Com49-2 | PKSP 4630 | Reverse | GGTCTCTAAGCCCTCCAGTGAGCTGCTTTATAA | This work |
| Com49-3 | PKSP 4631 | Forward | GGTCTCAAGGTAGATCAGTTTAGCGTGGAATGT | This work |
| Com49-3 | PKSP 4632 | Reverse | GGTCTCTAAGCCTCTCACTTCATGGTTCTTTGC | This work |
| Com49-4 | PKSP 4633 | Forward | GGTCTCAAGGTAGAAGCAGTACTTTCAGTCCTC | This work |
| Com49-4 | PKSP 4634 | Reverse | GGTCTCTAAGCTGAATTGTTGATAGTGTGCTCTG | This work |
| Com49-5 | PKSP 4635 | Forward | GGTCTCAAGGTTGGTTGGAGAAGCATGGGG | This work |
| Com49-5 | PKSP 4636 | Reverse | GGTCTCTAAGCAAGATTGAACTCCTCCTGCCC | This work |
| Com49-6 | PKSP 4637 | Forward | GGTCTCAAGGTTGCAAGACTTCCCAACCTTG | This work |
| Com49-6 | PKSP 4638 | Reverse | GGTCTCTAAGCACTACATCGTCTAGTTTCAACAGTT | This work |
| *NbPtr1* | PKSP 4665 | Forward | GGTCTCAAGGTATGGCAGAATTTTTCTTGTTCAAC | This work |
| *NbPtr1* | PKSP 4666 | Reverse | GGTCTCGAAGCCTTCTTCTTAAGGCTTTCCTGAAA | This work |
| *NbZAR1* | PKSP 3464 | Forward | ATTCGAATTCCTCACTACAGTTGAGCTATGATGAA | This work |

| | | | | |
|---|---|---|---|---|
| *NbZAR1* | PKSP 3465 | Reverse | TAGAGGATCCGCAATGTCGGAATGGATTTTGTT | This work |
| *NbZAR1* | PKSP 4354 | Forward | GGTCTCCAGGTACTCACTACAGTTGAGCTATGATGAA | This work |
| *NbZAR1* | PKSP 4355 | Reverse | GGTCTCCAAGCTGCAATGTCGGAATGGATTTTGTT | This work |
| *CaPtr1* | PKSP 4609 | Forward | GGTCTCAAGGTATGGCGGAATCGTTCTTGTT | This work |
| *CaPtr1* | PKSP 4610 | Reverse | GGTCTCTAAGCCTTCTTCCCAAAGCTTTTCTGG | This work |
| *CaZAR1* | PKSP 4605 | Forward | GGTCTCAAGGTTCAAGAACGAACTGCTATTCATG | This work |
| *CaZAR1* | PKSP 4606 | Reverse | GGTCTCTAAGCTAAGTGGCACTCCAAGGTATG | This work |
| *NbRIN4-1* | PKSP 4160 | Forward | GGTCTCCGGAGCACAAACAGAGGAACCAATTGG | This work |
| *NbRIN4-1* | PKSP 4161 | Reverse | GGTCTCACTACTATTCTTTCCTTCCCAAGCAGG | This work |
| *NbRIN4-2* | PKSP 4162 | Forward | GGTCTCTGTAGATACAACACTCCCTATACTGTCTTTTTT | This work |
| *NbRIN4-2* | PKSP 4163 | Reverse | GGTCTCAAGATACTCGTCCTGATATCTTTGCCT | This work |
| *NbRIN4-3* | PKSP 4164 | Forward | GGTCTCTATCTCAAGTATAGAAGAGTATCCGAGTAGACC | This work |
| *NbRIN4-3* | PKSP 4165 | Reverse | GGTCTCTAAGCATCTTCCAGGTGTAGAGGGAG | This work |
| **Primers used for knocking out *hopQ1-1* from *P. synringae* pv. *tomato* DC3000** | | | | |
| HopQ1-1 flanking L | PKSP 3224 | Forward | GGTCTCAAATGGTGCCTCCTTTGATTATCGAGT | This work |
| HopQ1-1 flanking L | PKSP 3225 | Reverse | GGTCTCACGAACCCGTAGTGCCGACAAATG | This work |
| HopQ1-1 flanking R | PKSP 3226 | Forward | GGTCTCATTCGTGAGTCGATCTCCTGGATAGAT | This work |
| HopQ1-1 flanking R | PKSP 3227 | Reverse | GGTCTCAAAGCAGCGCTCAACCTGACCAG | This work |
| **Primers used for semi- or quantitative PCR** | | | | |
| *NbActin* | PKSP 4380 | Forward | GATGAAGATACTCACAGAAAGA | This work |
| *NbActin* | PKSP 4379 | Reverse | GTGGTTTCATGAATGCCAGCA | This work |
| *NbPtr1* | PKSP 4667 | Forward | CTCAATATTGGTAAAATGTCTGATTGT | This work |
| *NbPtr1* | PKSP 4777 | Reverse | GAAATGTGCATTTGTAGGGGAG | This work |
| *NbRIN4-1* | PKSP 2899 | Forward | GGTCTCAAATGGCACGTCCAAATGTTCC | This work |
| *NbRIN4-1* | PKSP 4358 | Reverse | CTGGAGGTGGGGGAGTAAGA | This work |
| *NbRIN4-2* | PKSP 4359 | Forward | TGCTTCGCCTGTTTGGGAAG | This work |
| *NbRIN4-2* | PKSP 4360 | Reverse | TCATTCTCATCCCATTCGCC | This work |
| *NbRIN4-3* | PKSP 4361 | Forward | CCAGGAGAGATGATGTAGAA | This work |
| *NbRIN4-3* | PKSP 4362 | Reverse | GACGTTCATGCCTTGCTGTT | This work |
| *TRV CP* | PKSP 3055 | Forward | ACGATTCTTGGGTGGAATCA | This work |

| | | | | |
|---|---|---|---|---|
| *TRV CP* | PKSP3056 | Reverse | CGGTGCAGATGAACTAGCAG | This work |
| **Additional primers used for creating constructs for agroinfiltration** | | | | |
| *avrB* | oCM275 | Forward | CACCATGGGCTGCGTCTCGTC | This work |
| *avrB* | oCM276 | Reverse | GAAAAAGCAATCAGAATCTAGCAAGC | This work |
| *avrRpm1* | oCM277 | Forward | CACCATGGGCTGTGTATCGAG | This work |
| *avrRpm1* | oCM278 | Reverse | AAAGTCATCTTCTGAGTCAGACTGAAC | This work |
| *hopZ5* | oCM279 | Forward | CACCATGGGACTTTGTGCATCAAAAC | This work |
| *hopZ5* | oCM280 | Reverse | GGATTCTATCGCTTTTCTTATTTTTCGTAGTCTCG | This work |

**Methods S1 RNA extraction and quantitative RT-PCR**

For total RNA extraction, 8 leaf discs of 5 mm diameter from five-week-old VIGSed *N. benthamiana* were sampled and frozen in liquid nitrogen. Total RNA was extracted using Trizol (Sigma-Aldrich, USA) reagent following the manufacturer's instructions. Subsequently, extracted RNA was treated with DNase, and 1 ug of RNA was used for cDNA synthesis using Toyobo ReverTraAce (Toyobo, Japan) kit. *N. benthamiana Actin* gene was used as a reference to compare the expression of VIGS target genes. Primers used in quantitative RT-PCR are listed in Table S3.

**Methods S2 Deletion of *hopQ1-1* from *Pseudomonas syringae* pv. *tomato* DC3000**

PCR products of LB and RB regions upstream (1.0-kb) and downstream (1.6-kb) of *hopQ1-1* gene were cloned into pUC19b vector and confirmed by sequencing (Table S3) (Jayaraman *et al.*, 2017). Then the PCR products were assembled in a Golden Gate compatible vector pK18-MobSacB (KanR) (Jayaraman *et al.*, 2020) and confirmed the correct assembly by a restriction enzyme test. The assembled construct was transformed into the target bacteria *Pseudomonas syringae* pv. *tomato* DC3000 using electroporation and plated on KB supplemented with Rif and Kan plates. Four transformant colonies were restreaked at 3-4 days post transformation on the new plates with the same selection. Two colonies from each transformant were inoculated in liquid KB culture with Kan selection, and each was spotted 100µL with a pipette on 5% sucrose LB plates and spread with a loop in a zig-zag fashion leaving heavy inoculum at the top of the plate and little at the bottom. Eight revertant colonies were restreaked per Kan-resistant transformant at 3-4 days post transformation on 10% sucrose LB plates. Then eight revertant colonies per transformant were screened by streaking them on LB plates with (no growth) or without (good growth) kanamycin. Finally, the presence of knockout band (primer F of LB and R of RB) and absence of WT *hopQ1-1* were confirmed by PCR (Table S3).

**Methods S3 AvrBsT complementation of *Xanthomonas perforans* 4B *ΔxopQ ΔavrBsT***

The *avrBsT* gene was introduced into *X. perforans* 4B *∆xopQ ∆avrBsT* (Schwartz et al., 2015) by triparental mating with modifications (Kvitko & Collmer, 2011; Kraus *et al.*, 2017) using *E. coli* Stellar carrying the helper plasmid pRK600. Briefly, the *X. perforans* 4B *∆xopQ ∆avrBsT* strain was grown on KB medium plates supplemented with glutamine at 16 g/liter for 2 days at 30°C to reduce the production of exopolysaccharides that might interfere with the mating process (Martin *et al.*, 1988). On the day before the conjugation, *E. coli* DH5α containing the donor plasmid and *E. coli* Stellar carrying the helper plasmid pRK600 were inoculated into liquid LB and grown overnight at 37 ºC. The cultures were subcultured in the morning of the conjugation until they reached an $OD_{600}$ of 0.5. The *X. perforans* 4B Δ*xopQ* Δ*avrBst* strain was resuspended in liquid KB media to a final $OD_{600}$ of 0.5. The subcultured *E. coli* strains were washed and brought to the same concentration in liquid KB. All strains were mixed 1:1:1, and the resulting solution was plated on a sterile nitrocellulose square on KB medium plates supplemented with glutamine. Plates were incubated for 1 day at 30°C, and then bacteria grown on the nitrocellulose squares were resuspended in 200 ml of KB media and plated on minimal mannitol-glutamate (MG) medium (mannitol at 10 g/liter, L- glutamic acid at 2 g/liter, $KH_2PO_4$ at 0.5 g/liter, NaCl at 0.2 g/liter, and $MgSO_4$ at 0.2 g/liter; final pH 7) supplemented with kanamycin (Kan) at 35 ug/ ml plates and incubated for 3 to 6 days at 30°C (Bronstein et al., 2008). The resulting transformants were selected via PCR of *avrBsT* and *xopJ4*.

**Methods S4 Generation of *Nicotiana benthamiana ptr1* mutant using CRISPR/Cas9**

To mutate the *NbPtr1* gene in *N. benthamiana* accession Nb-1 (Bombarely *et al.*, 2012), we designed 2 guide RNAs (NbPtr1a/b-1: gACAAGTATGTTCAGTAAAGTGG and NbPtr1a/b-2: gCAAACACATTCCTTTGTAAGGG) targeting the first exon of *NbPtr1* using the software Geneious R11 (Kearse *et al.*, 2012). Each gRNA cassette was cloned into a Cas9-expressing binary vector (p201N:Cas9) by Gibson assembly (Jacobs *et al.*, 2017). *N. benthamiana* transformation was performed at the biotechnology facility at the Boyce Thompson Institute. *Agrobacterium* cells containing each gRNA/Cas9 construct were pooled together and used for transformation into the *N. benthamiana*. To determine the mutation type, genomic DNA was extracted from the leaves of each transgenic plant using a modified CTAB method (Murray & Thompson, 1980). Genomic regions spanning the target site of the *NbPtr1* gene were amplified with specific primers (NbPtr1a_F: GTAAATGAGATCAGTTTAGCGTGGAATG and NbPtr1a_R: CATCCAGTACAAGTAAATACCTTTTCG) and sequenced at the Biotechnology Resource Center (BRC) at Cornell University. Geneious R11 and the web-tool called Tracking of Indels by Decomposition (TIDE; https://tide.deskgen.com) (Brinkman *et al.*, 2014) were used to determine the mutation type and frequency using the sequencing files (ab1. format) as described (Zhang *et al.*, 2020).

**Methods S5 *P. syringae* pv. *tomato* ∆*hopQ1-1* ∆*avrPto* ∆*avrPtoB* culture and transformation**

*Pseudomonas syringae* pv. tomato strains DC3000 ∆*hopQ1-1* ∆*avrPto* ∆*avrPtoB* (Kvitko *et al.*, 2009) were grown on King´s B (KB) semi-selective media at 30°C. Plasmids pCPP5372 (Oh *et al.*, 2007) carrying AvrRpt2, AvrB, AvrRpm1, and HopZ5 were introduced into DC3000 ∆*hopQ1-1* ∆*avrPto* ∆*avrPtoB* by electroporation. All *P. syringae* pv. tomato strains were stored in 20% glycerol + 60 mM sucrose at –80°C.

**Methods S6 *P. syringae* pv. *tomato* inoculation and population assays in tomato**

*P. syringae* pv. tomato DC3000 ∆*hopQ1-1*∆*avrPto*∆*avrPtoB* expressing AvrRpt2, AvrB, AvrRpm1, HopZ5, and the empty vector were grown on KB plates for 2 days at 30°C. Strains were diluted in 10 mM $MgCl_2$ + 0.002% Silwet L-77 at a final concentration of 2 x $10^4$ CFU $ml^{-1}$. Four-week-old LA4245-R and LA4245-S plants were vacuum infiltrated, and three leaf disk samples (7 mm in diameter) were collected 2 days post inoculation (dpi) to quantify bacterial populations. The experiments were repeated three times. The results shown are the mean of three independent experiments using three biological replicates per strain, including the standard error of the mean. Statistical analyses were performed using Prism 6.0 (GraphPad Software).
